# Ecological and functional niches comparison reveals differentiated resource-use strategies and ecological thresholds in four key floating-leaved macrophytes

**DOI:** 10.1101/2024.06.13.598787

**Authors:** Alice Dalla Vecchia, Maria Beatrice Castellani, Mattia M. Azzella, Rossano Bolpagni

## Abstract

Niche theory has been widely used in ecology; however, few studies have attempted to combine information on functional and ecological niches (i.e. variation in traits and environmental requirements), especially for freshwater macrophytes.

In this study we aim to describe the functional and ecological niches of four key nymphaeid species (*Nuphar lutea*, *Nymphaea alba*, *Nelumbo nucifera* and *Nymphoides peltata*) to investigate their environmental tolerance and functional adaptability.

Twelve Italian populations per species were sampled. Functional and ecological niches were determined using hypervolumes based on eight functional traits and environmental variables, related to leaf structure and economics spectrum, and to water and sediment quality.

Among the three Italian native species, *N. lutea* and *N. alba* showed intermediate niche size and position in the functional and ecological space, although *N. alba* appears to be less competitive due to a small functional niche and lower trait performance, which could explain its general tendency to decline. *N. peltata* appeared more specialized in its environmental requirements and characterized by highly acquisitive leaves, while the invasive alien *N. nucifera* exerted its competitive success by distinguishing its functional niche and expanding its ecological niche, through high investment of resources in leaves.

Overall, all four target species share similar ecological niches, colonizing eutrophic ecosystems typical of intensive agricultural landscapes, but show different patterns in their functional niche. We demonstrate the applicability of an approach based on both functional and ecological niches to unravel species’ adaptation and strategies.

## 1. INTRODUCTION

Niche theory has long been used in ecology studies (Devictor et al. 2010). Definitions of niche are varied, but generally refer to the way a species is able to use and adapt to the resources available in its habitat, describing the ecological niche (Grinnel 1917), or to the effect of a species in the environment, considering its characteristics as a functional niche (Elton 1927; Devictor et al. 2010). The ecological niche is by far the most studied in vegetation science: indeed, it can give valuable information highlighting the behavior of generalist vs. specialized species (Devictor et al. 2010), and reflects the effect of habitat filtering (Viana et al. 2016). The size and conditions that describe niches can cause important implications for species distribution, requirements and conservation concerns in a changing environment (Papuga et al. 2018; Liu et al. 2021), and potentially help understanding the dynamics of (invasive) species spread (Montagnani et al. 2022; Robichaud & Rooney 2022). Indeed, functional traits – that are any morphological, physiological or phenological feature measurable at the individual level and affecting plant fitness (McGill et al. 2006) – can substantially implement the niche concept by defining the extent of species responses (Kearney et al. 2010; Ajal et al. 2022). Leaf structural, chemical and physiological traits, among others, can reveal important trade-offs in plant resource-use strategies, which are closely linked to ecological conditions (Wright et al. 2004).

Macrophytes exhibit extremely wide traits variation in response to aquatic habitat peculiarities and steep ecological gradients (Pierce et al. 2012; Dalla Vecchia & Bolpagni 2022). In these plants, phenotypic plasticity interacts with environmental tolerance, greatly influencing their ability to persist under different conditions (De Wilde et al. 2014). Indeed, including intraspecific trait variability in ecological studies can capture more information that can better describe the tolerance of these species (Puglielli et al. 2024). On one hand, macrophytes are facing a rapid and dramatic decline worldwide due to direct anthropic pressure, climate change and habitat degradation (O’Hare et al. 2018). On the other side, however, many macrophyte species are spreading worldwide and represent some of the most impactful invasive species, causing negative effects on ecosystems and nuisance to human activities (Strayer 2010; Hussner et al. 2017; Bolpagni 2021). Therefore, a combined approach, integrating ecological and functional niche, can lead to a more complete understanding of the extent of variation in resource-use strategies and functional adaptation of macrophytes (Lukács et al. 2017), allowing us to catch possible mechanistic explanations for their decline, migration or invasive behavior.

Nymphaeids (water lilies) are aquatic rooted plants with rosette-shaped leaves that are mostly floating or emergent, reaching the water surface with elongated petioles. Leaves can vary in size (Pierce et al. 2012), and can entirely cover the colonized water bodies, a strategy that helps them outcompete other macrophyte growth forms and results in mostly monospecific populations (Paillisson & Marion 2011; Nowak et al. 2015). Nymphaeids are a key component of aquatic ecosystems, influencing sediments, water and atmosphere, but have so far received little attention compared to submerged vegetation in ecological studies (Klok & van der Velde 2017). Nevertheless, some of these macrophytes are showing declining trends, such as *Nymphaea* spp., although a full understanding of the responses of these species to changing environmental conditions has not yet been achieved (Parveen et al. 2022).

Invasive nymphaeid species are also forcing native species into severe regression as they rapidly colonize water bodies: for example, the invasive *Nelumbo nucifera* Gaertn. is replacing native stands of *Nymphaea alba* L. and *Nuphar lutea* (L.) Sm. in shallow lakes of Northern Italy (Villa et al. 2018; Pinardi et al. 2021), even though *N. lutea* is considered a top competitor species (Temmink et al. 2021). Indeed, nymphaeids also include invasive species that are detrimental to ecosystem functioning and practical human utility (Darbyshire & Francis 2008).

Based on a CSR functional classification, which describes species as Competitive, Stress-tolerant or Ruderal (Grime & Pierce 2012), Dalle Fratte et al. (2019) did not observe a difference between native and alien species of nymphaeids in their strategies, all clustering together in the most competitive edge of the spectrum compared to other aquatic or terrestrial plants. Overall, it remains unclear whether the invasive species share the same niche as the natives, outcompeting them, or whether they are able to occupy a new niche, left unexploited by the native species (Loiola et al. 2018; Dalle Fratte et al. 2019).

It is hence of great interest to evaluate the degree of niche uniqueness – namely, the proportion of a species’ niche not shared with others – to quantify differences among species and catch functional or ecological specialization. Previous studies on nymphaeids have shown great variability in traits in response to environmental (trophic) gradients (Dalla Vecchia & Bolpagni 2022; Castellani et al. 2023; Dalla Vecchia et al. 2024). The contrasting results regarding their performance and resource-use strategies under different eutrophic conditions stress the need to combine functional and ecological information (Pełechaty 2007).

In this study we aim at evaluating the size and uniqueness of the ecological and functional niches (defined here as a multidimensional space given by the set of environmental conditions in which a species lives and the variability of its functional traits, respectively) of four key nymphaeid species, and to compare them on the basis of information about both niches. We ultimately aimed to interpret the differences in niches among species, their links to inform about leaves resource-use strategies of each species and associated ecosystem impacts. We focus on *N. lutea*, *N. alba*, *Nymphoides peltata* (S.G.Gmel.) Kuntze and *N. nucifera*, four key nymphaeid plants with varying degree of rarity (*N. peltata* > *N. alba* >> *N. nucifera* ∼ *N. lutea*) and competitive behaviour (*N. nucifera* > *N. lutea >> N. alba* ∼ *N. peltata*). We hypothesize that i) the species showing a stronger tendency to decline, namely *N. alba* and *N. peltata*, will show smaller niche sizes and ii) greater niche overlap with other species, indicative of lower environmental tolerance and traits adaptability. We also hypothesize that iii) the more competitive species, such as *N. nucifera* and *N. lutea* will show large niche sizes, and that iv) their functional niche would highly overlap with other native species (especially for *N. nucifera*), suggesting, on the other hand, broad tolerance and adaptation to a range of environmental conditions.

## 2. MATERIALS AND METHODS

### 2.1 Study area

Plants and related environmental descriptors were sampled at several locations in Italy (Fig. 1) between 2020 and 2022, where the presence of the species of interest was known from expert-based observation, previous studies (e.g., Lastrucci et al. 2014; Pinardi et al. 2021) or Italian websites with updated records (www.actaplantarum.org). Study sites included 11 lakes, among which Cei Lake (Trento, 45.9° N 11.0° E), Varese Lake (Varese, 45.8° N 8.7° E), Comabbio Lake (Varese, 45.8° N 8.7° E), Pusiano Lake (Como, 45.8° N 9.3° E), Annone Lake (Lecco, 45.8° N 9.4° E) and Fimon Lake (Vicenza, 45.5° N 11.5° E) are sub-alpine lakes, ranging in size from 0.045 km^2^ (Cei Lake) to 14.5 km^2^ (Varese Lake). Other lakes are: the fluvial lakes system of Mantua (45.2° N 10.7° E); Gorro Lake located in the Northern Apennines (Parma, 44.5° N 9.9° E); Chiusi Lake, a shallow lake remnant of ancient wetlands (Siena, 43.1° N 12.0° E); the Monticchio Grande Lake (Potenza, 40.9° N 15.6° E), a small shallow volcanic lake in southern Italy. Samples were collected also from two wetlands, the Torbiere del Sebino, south of Iseo Lake (Brescia, 45.6° N 10.0° E) and the Vallazza, south of the Mantua lakes system (Mantua, 45.1° N 10.8° E), and from a pond in Massarosa (Lucca, 43.9° N 10.3° E). Lastly, we included in the study some artificial large canals along the last stretch of the Po River (in the province of Ferrara, from 45.0° N 11.9° E eastwards). They show minimal to no water flow (< 0.2 m s^−1^) and can therefore be ecologically attributed to stagnant environments. Altitude of sampling sites varied from 3 m a.s.l. (Codigoro, Ferrara) to 920 m a.s.l. (Cei Lake, Trento). A summary table of environmental conditions of each lake can be found in Supplementary Information, Heading 1.

**Figure 1.**
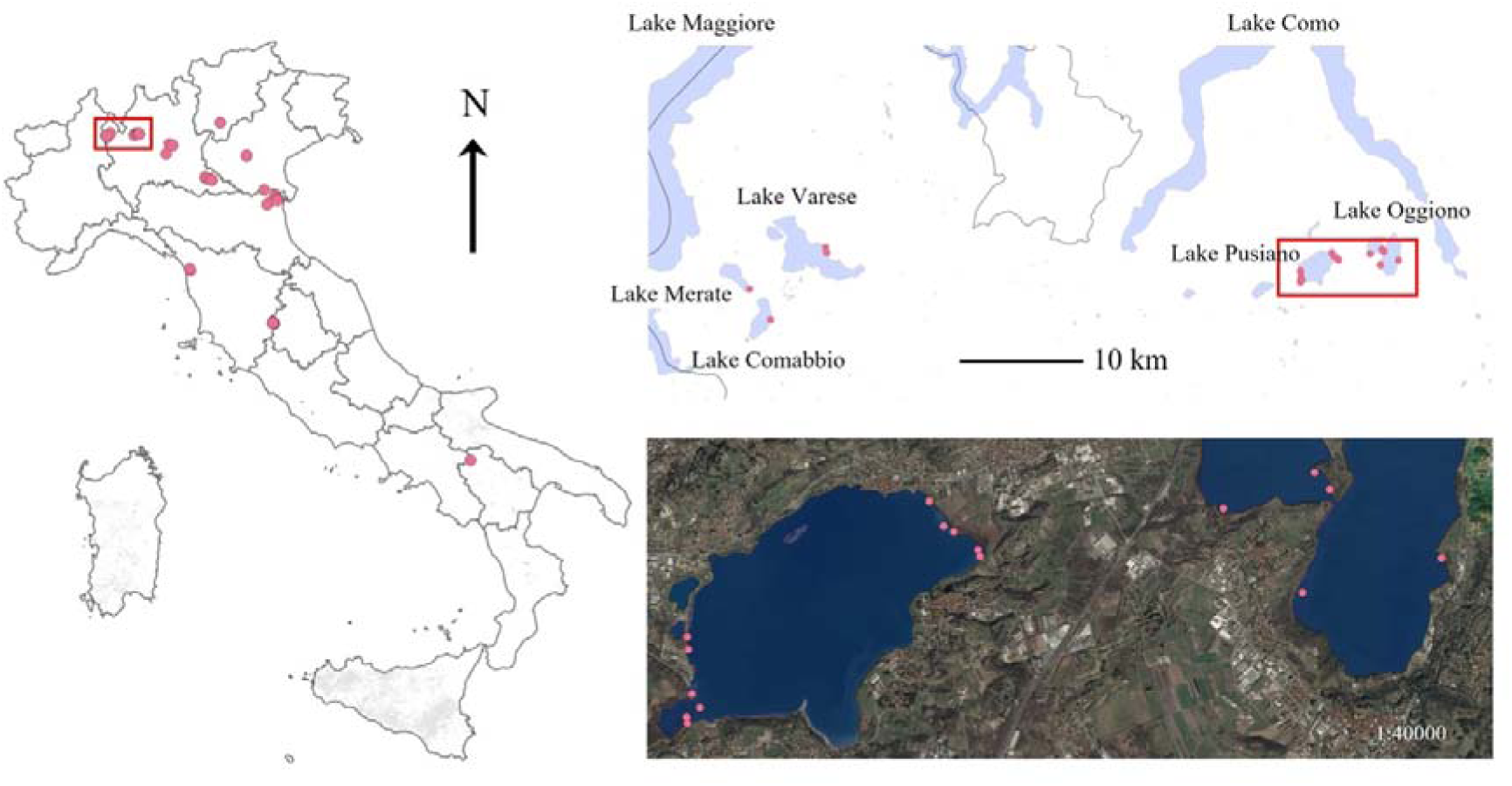
Map of the study sites located in Italy, and a focus on the alpine lakes near Lake Maggiore and Lake Como sampling area, showing the spatial arrangement of sampling plots.

### 2.2 Ecological niche sampling

Water physical and chemical, and sediment parameters were characterized for each population. For the purpose of this study, we use the term “population” to refer to each sampled macrophyte stand, defined by a 4×4 m^2^ plot and isolated from other macrophytes stands of the same species by at least 200 m, with no presence between them. Depending on the extent and heterogeneity of the study sites, some lakes and canals could host more than one population. *In situ*, water temperature, electrical conductivity, pH and dissolved oxygen were measured with a multiparameter probe (Eureka Manta2), as well as water depth. A sample of water was collected, immediately filtered with 0.7 µm pore size glass fiber filters (Whatman) and kept refrigerated at 4° C until processing. Dissolved inorganic carbon was determined by titration with 0.1 N HCl (Anderson et al. 1986), and soluble reactive phosphorus (detection limit 4.00 μg L^−1^) was determined spectrophotometrically following Valderrama (1977). Both analyses were carried out within 48 hours from sampling. The remaining sample was frozen for further analyses. Dissolved reactive silica was then determined spectrophotometrically (Lienig et al. 1978). After filtering the samples with 0.2 µm pore size nylon filters, dissolved anions and cations were determined by ion chromatography (883 Basic IC plus Metrohm, Herisau, Switzerland). A 50 mL sediment sample was collected from the upper 5 cm sediment layer in each population, stored in a falcon tube and frozen until processing. In the laboratory, it was homogenized, and a subsample of 5 mL was used for determination of sediment density, porosity and water content by weighing the fresh and dry weight after drying in the oven at 50°C until constant weight. Around 0.3 g of dry sediment were used for the determination of organic matter content after combustion at 450° C for 4 h, using the gravimetric method (Buchanan et al. 1984). After acid extraction of a subsample of the incinerated sediment, total phosphorus, including organic and inorganic phosphorus, was analyzed spectrophotometrically (Aspila et al. 1976).

### 2.3 Species and functional niche sampling

We investigated four species of nymphaeids: *N. nucifera*, *N. lutea*, *N. alba* and *N. peltata*. *N. nucifera* is an invasive species in Italy, introduced in the 1920s (Tóth et al. 2019) and forms dense populations with floating and emergent leaves, while the latter three are native species. All of them are rooted floating-leaved species. For each species, 12 populations were characterized. We chose to measure leaf traits related to the leaf economics spectrum because they reflect the plants resource-use strategies (Wright et al. 2004), especially in nymphaeids, where leaves represent most of the above-ground biomass (Brock et al. 1983a). Besides, the use of leaf economics spectrum traits is widely consolidated in the literature, which would allow our results to be more comparable with previous studies (Pierce et al. 2012; Pan et al. 2019). Finally, variability of these traits in response to variation in water and sediment parameters was already observed in *N. lutea* (Dalla Vecchia & Bolpagni 2022; Dalla Vecchia et al. 2024). In each population, eight young, i.e., fully expanded leaves with no sign of damage, or illness, or herbivory were collected from eight distinct plants, paying attention to collect the full petiole length. The leaves were immediately refrigerated in airtight plastic bags to avoid dehydration. Fresh weight as well as leaf area were determined on five leaves within hours from collection, using a precision scale and a scanner at 300 dpi resolution. Leaf area was successively determined using the software imageJ (Rasband 1997-2018). At the same time, leaf disks of known area (24.8 mm^2^) were cut out of fresh blades from the remaining three leaves and frozen in airtight plastic containers for further pigments analyses. Weighed leaves were then dried in the oven at 50° C until constant weight, to determine dry weight. Leaf fresh weight, leaf dry weight and leaf area were measured on blades and petioles separately. We then calculated leaf dry matter content and specific leaf area following Pérez-Harguindeguy et al. (2013). We also calculated two petiole-related traits: proportion of petiole area and proportion of petiole dry weight, measured as the ratio between the petiole area or dry weight and the total leaf area or dry weight, respectively. These two traits indicate the resource investment of plants in the structural support of leaves and colonization of water surface. Dry material of leaf blades was pooled together, ground to fine powder with liquid nitrogen and stored for elemental composition analyses (phosphorus, nitrogen and carbon). From the pooled material, three aliquots were used for phosphorus content determination, measured spectrophotometrically after acid extraction as described for the sediments, and one more aliquot was used for carbon and nitrogen determination by combustion analysis with an elemental analyzer (Thermo FlashEA 1112). Leaf chlorophyll-*a*, chlorophyll-*b* and carotenoids content were determined on a fresh weight and area basis after extraction for 24 h in 80% acetone, and the solution was read spectrophotometrically according to Wellburn (1994). Ratios between chlorophyll-*a* and chlorophyll-*b*, and between chlorophylls and carotenoids were subsequently calculated.

### 2.4 Statistical analyses

All analyses were carried out in R environment (R Core Team 2022), and graphical representations of all results were created with packages *ggplot2* (Wickham 2016), *ggbiplot* (Vu 2011) and *rgl* (Murdoch & Adler 2023).

To summarize the ecological and functional information useful for the computation of the niches, two principal component analyses (PCA) were carried out, one on the environmental parameters and one on traits. The number of variables included in the PCAs was reduced removing redundant variables, defined by a Pearson’s correlation coefficient >|0.7| (see Supplementary Information, Heading 2 for the results of the correlation analysis). All variables included were transformed to have mean of zero and unit variance before running the PCA. The function `ggscreeplot` from the package *ggplot2* (Wickham 2016) was used to determine the optimal number of axes to be included in the niche determination. The package *hypervolume* (Blonder et al. 2022) was implemented and integrated to quantify the size and uniqueness of the niche. The niche size, i.e., the volume occupied by a species relative to the volume occupied by all species in the dataset, was used to compare the investigated species. Hypervolumes were built with the gaussian kernel method, being the most appropriate for functional data and fundamental niche application (Blonder et al. 2018). This method uses elliptic random sampling to create clouds of data points around the observed data points, assuming that sampled data points within the hypervolume would be close to observed data points (Blonder et al. 2018). However, the number of observations influences the final hypervolume size, because higher uncertainty (i.e., fewer data points) results in bigger hypervolume size. To perform set operations, we subset hypervolumes of each species to the same random density using the function `hypervolume_n_occupancy`. Mean absolute error (MAE) and Root mean squared error (RMSE) were used to evaluate the accuracy of the subset hypervolumes (Laini et al. 2023). The relative niche size of each species was then calculated as its absolute volume divided by the volume of the union of input hypervolumes. Relative niche uniqueness, on the other hand, was calculated as the absolute niche uniqueness of a species divided by its absolute niche size. Niche uniqueness also reflects niche overlap, as “1 - relative uniqueness” = relative overlap (i.e., proportion of a species niche that is shared with other species). To obtain uncertainty measures of niche size and niche overlap, bootstrapping was implemented on 199 permutations (to optimize accuracy of results and computation performance) of each species niche, which were then used to calculate 199 relative total occupancies. Pairwise niche size comparison among species, as well as niche uniqueness comparison, were tested setting the significance level at 0.05. We also obtained estimates of pairwise niche intersection, which is the proportion of niche overlapping between two species compared to the union of volumes of all species.

To assess the significance of differences in environmental conditions and functional traits among the four investigated species we used linear mixed models, followed by post-hoc pairwise comparisons tests. In the linear mixed models, the first three PCA axes of both functional and ecological niches were tested against species (fixed effect factor), year of sampling and population identity (random effect factors). Normality of residuals was assessed visually to meet linear mixed model assumptions. The analysis was carried out using the packages *lme4* (Bates et al. 2015) and post-hoc comparisons using *emmeans* (Lenth 2022). Moreover, differences among species in single traits and environmental variables were tested similarly using linear mixed models as well as ANOVA and Kruskal Wallis non-parametric tests when residuals of environmental variables were not homogeneously distributed (Results are reported in Supplementary Information, Heading 3).

## 3. RESULTS

Based on Pearson’s correlation analysis, the niche analyses included eight environmental variables – depth, pH, electrical conductivity, dissolved inorganic carbon, nitrate, dissolved reactive silica, sediment organic matter content and sediment phosphorus content – and eight functional traits – leaf area, proportion of petiole dry weight, leaf dry matter content, specific leaf area, total chlorophylls, chlorophyll-a/chlorophyll-b, phosphorus and carbon content (Table 1). Phosphate and sediment density were excluded as they were highly correlated with electrical conductivity (P=0.72) and sediment organic matter content (P=-0.74), respectively. As for the functional traits, leaf size traits (area, fresh and dry weight, proportion of petiole area) were highly positively correlated (Supplementary Information, Heading 2). Leaf area was also positively correlated with pigments content per fresh weight (P>0.83), so we selected leaf area and total chlorophylls per surface for the PCA analysis. Leaf nitrogen content was highly correlated with leaf phosphorus content (P=0.75), and we selected the latter.

**Table 1:**
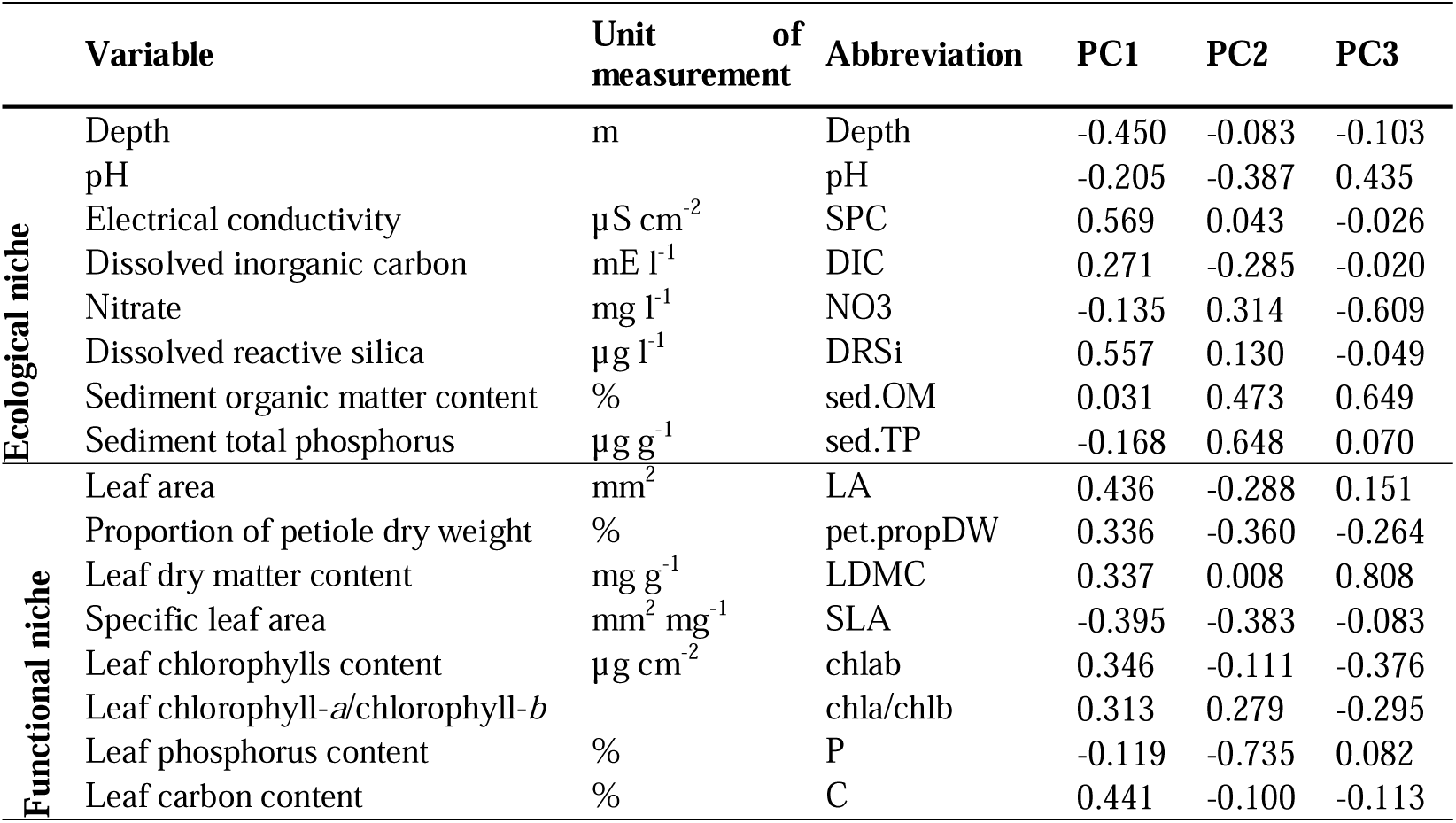
List of environmental variables and functional traits included in the niches’ computation. Loadings of the axes used in the analyses are reported.

|  | Variable | Unit of measurement | Abbreviation | PC1 | PC2 | PC3 |
| --- | --- | --- | --- | --- | --- | --- |
| Ecological niche | Depth | m | Depth | -0.450 | -0.083 | -0.103 |
|  | pH |  | pH | -0.205 | -0.387 | 0.435 |
| | Electrical conductivity | $\mu\text{S cm}^{-2}$ | SPC | 0.569 | 0.043 | -0.026 |
| | Dissolved inorganic carbon | $\text{mE l}^{-1}$ | DIC | 0.271 | -0.285 | -0.020 |
| | Nitrate | $\text{mg l}^{-1}$ | NO3 | -0.135 | 0.314 | -0.609 |
| | Dissolved reactive silica | $\mu\text{g l}^{-1}$ | DRSi | 0.557 | 0.130 | -0.049 |
|  | Sediment organic matter content | % | sed.OM | 0.031 | 0.473 | 0.649 |
| | Sediment total phosphorus | $\mu\text{g g}^{-1}$ | sed.TP | -0.168 | 0.648 | 0.070 |
| Functional niche | Leaf area | $\text{mm}^2$ | LA | 0.436 | -0.288 | 0.151 |
|  | Proportion of petiole dry weight | % | pet.propDW | 0.336 | -0.360 | -0.264 |
| | Leaf dry matter content | $\text{mg g}^{-1}$ | LDMC | 0.337 | 0.008 | 0.808 |
| | Specific leaf area | $\text{mm}^2 \text{mg}^{-1}$ | SLA | -0.395 | -0.383 | -0.083 |
| | Leaf chlorophylls content | $\mu\text{g cm}^{-2}$ | chlab | 0.346 | -0.111 | -0.376 |
|  | Leaf chlorophyll-a/chlorophyll-b |  | chla/chlb | 0.313 | 0.279 | -0.295 |
|  | Leaf phosphorus content | % | P | -0.119 | -0.735 | 0.082 |
|  | Leaf carbon content | % | C | 0.441 | -0.100 | -0.113 |

### 3.1 Ecological niches

The first three PCA axes were used in the hypervolumes computation, and together explained 60.0 % of the variance in the data (Fig. 2A-B-C). The first axis (25.4% explained variance) was related to water nutrients at the positive end (electrical conductivity, highly correlated with phosphate, and dissolved reactive silica) and with water depth at the negative end (Table 1). The second axis (20.1% explained variance) was more related to sediment features, with sediment phosphorus and organic matter content (negatively correlated to sediment density) associated with positive axis values. The third axis (14.4% explained variance) was related with both water and sediment quality, with sediment organic matter content and pH at the positive edge and nitrate at the negative edge of the axis. *N. nucifera* showed the highest mean relative niche size (75.9%), followed by *N. lutea* (24.4%), *N. alba* (14.4%) and *N. peltata* (5.6%). However, high uncertainty was associated to the ecological niche volume and no significant differences were observed among any species based on bootstrap results with 95% confidence intervals (Fig. 3A, Supporting Information, Heading 4).

**Figure 2.**
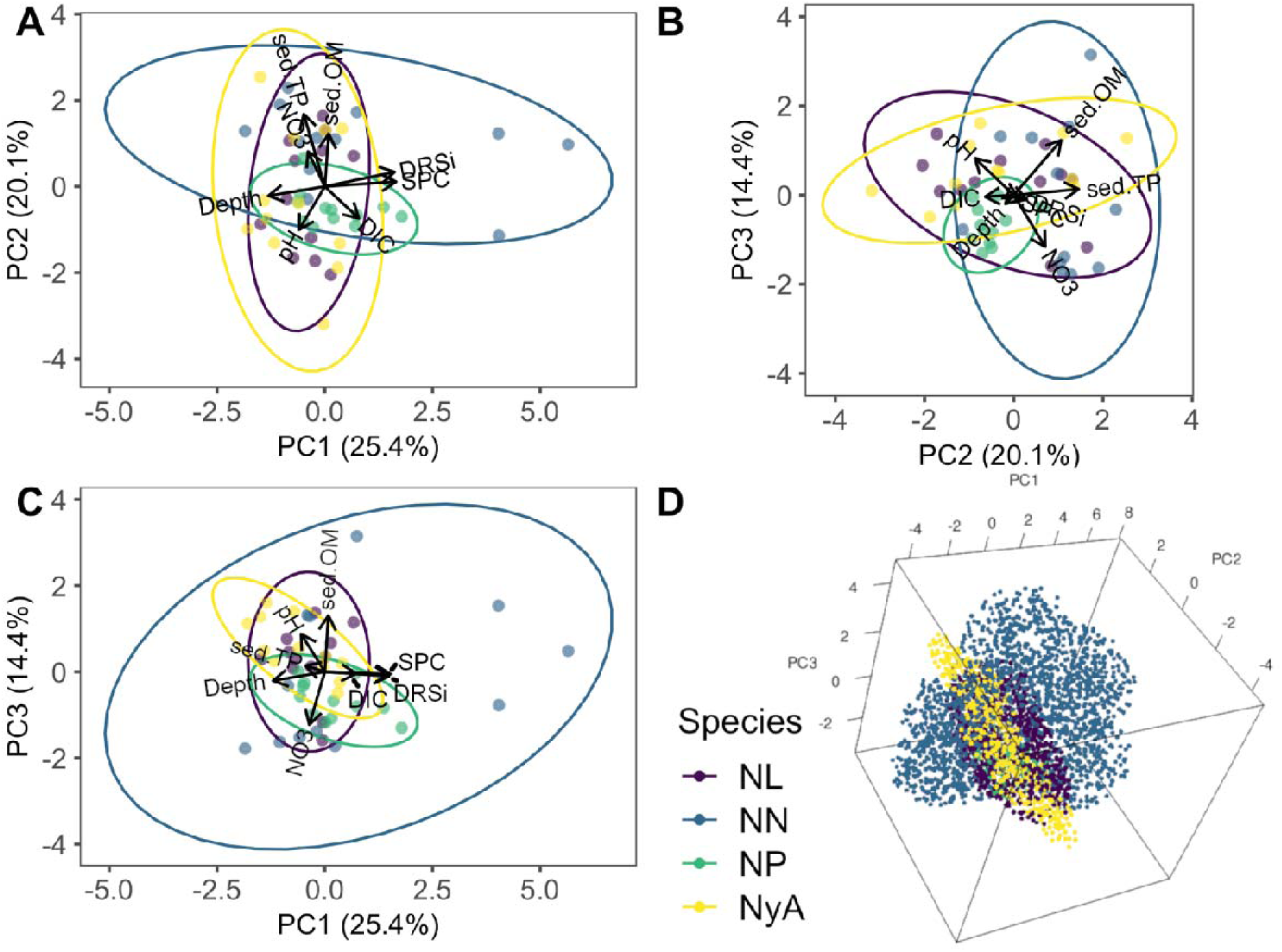
First three axes of environmental PCA (A-B-C) and 3-dimensional representation of the ecological niche (D). Percentage of explained variance of each axis is indicated in brackets. NL = *N. lutea*, NN = *N. nucifera*, NP = *N. peltata*, NyA = *N. alba*. For full environmental variables names see Table 1.

**Figure 3.**
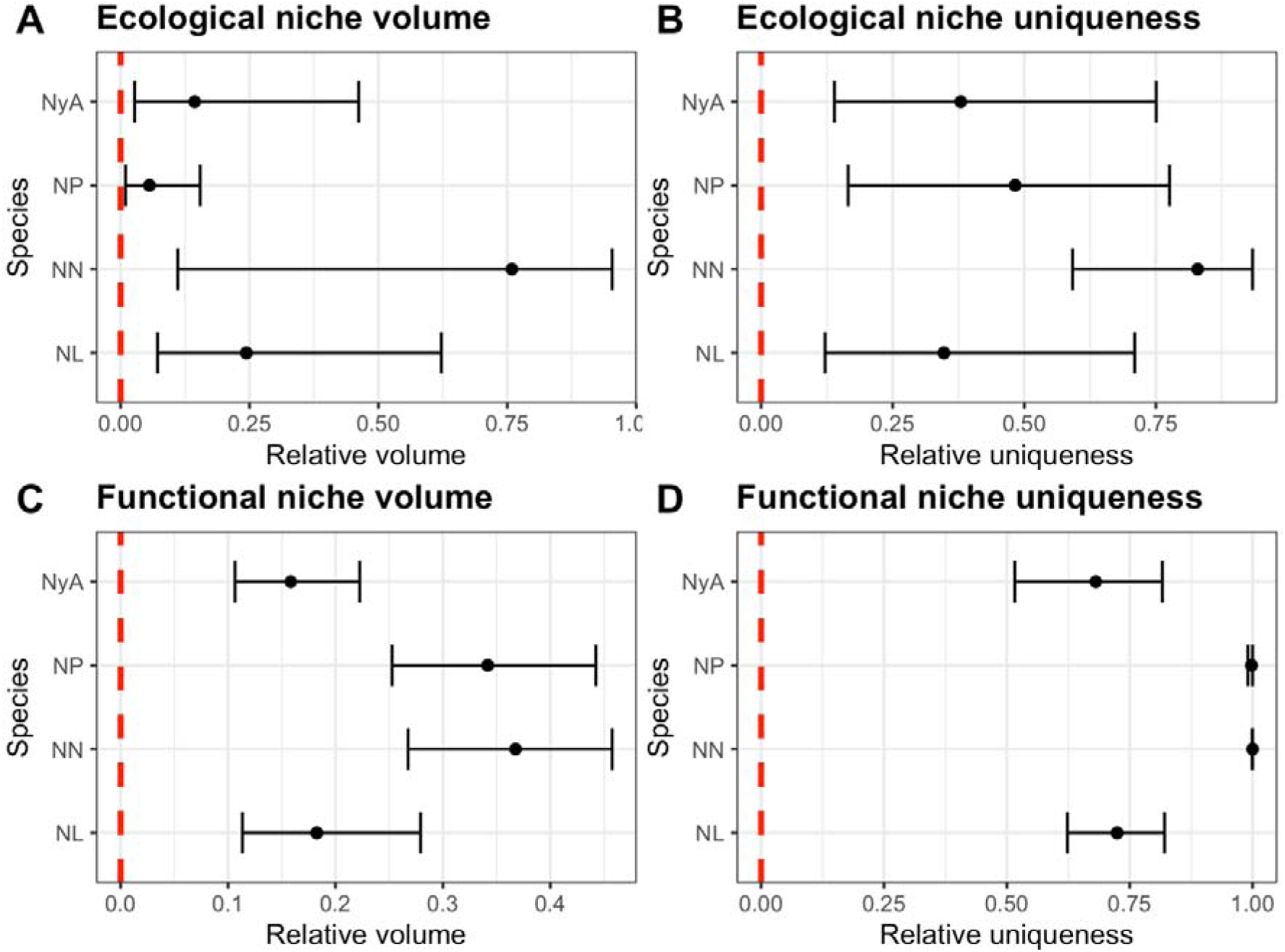
Uncertainty estimates of hypervolumes size (A-C) and uniqueness (B-D), obtained with bootstrapping on 199 permutated hypervolumes on each species. Relative niche size is the proportion of niche occupied by each species relative to the total niche occupied by all species. Relative niche uniqueness is the proportion of unique niche of each species relative to its total niche volume. Significant level was set at 0.05. Mean differences, 2.5 and 97.5 quantiles are provided in Supporting Information, Heading 4. Significant differences between species occur if bars do not overlap. NL=*N. lutea*, NN=*N. nucifera*, NP=*N. peltata*, NyA=*N. alba*.

*N. nucifera* showed the highest relative niche uniqueness, with 82.9% of its ecological niche, on average, not shared with any other species, followed by *N. peltata* (48.3%), *N. alba* (37.9%) and *N. lutea* (34.7%). In this case too, no species showed significantly different values of niche uniqueness based on bootstrap results (Fig. 3B, Supporting information, Heading 4). *N. lutea* presented a significant niche overlap with all other species, which was highest with *N. alba* (average 17.8% of all species volumes, Supplementary Information, Heading 4).

The results obtained suggest that all the investigated nymphaeid species share similar ecological conditions, though *N. peltata* presents a relatively smaller niche size (though the difference is not significant), due to its preference of sites with lower sediment organic matter content and higher sediment density. *N. nucifera*, on the other hand, showed a bigger and more unique niche (Fig. 2D), mostly due to three of its populations found at sites with very high electrical conductivity (>1300 µS cm^−1^). This was corroborated by the linear mixed models that highlighted no significant difference among species along PC1 (Fig. 4A). *N. nucifera* had significantly higher loadings of PC2 compared to *N. peltata* and *N. alba*, indicating a preference of the former species for sites with higher sediment total phosphorus and organic matter content (Fig. 4B, Supplementary Information, Heading 5). *N. peltata*, instead, showed significantly lower loadings of PC3 than *N. alba*, highlighting its presence in sites with higher sediment density and nitrate concentration (Fig. 4C).

**Figure 4.**
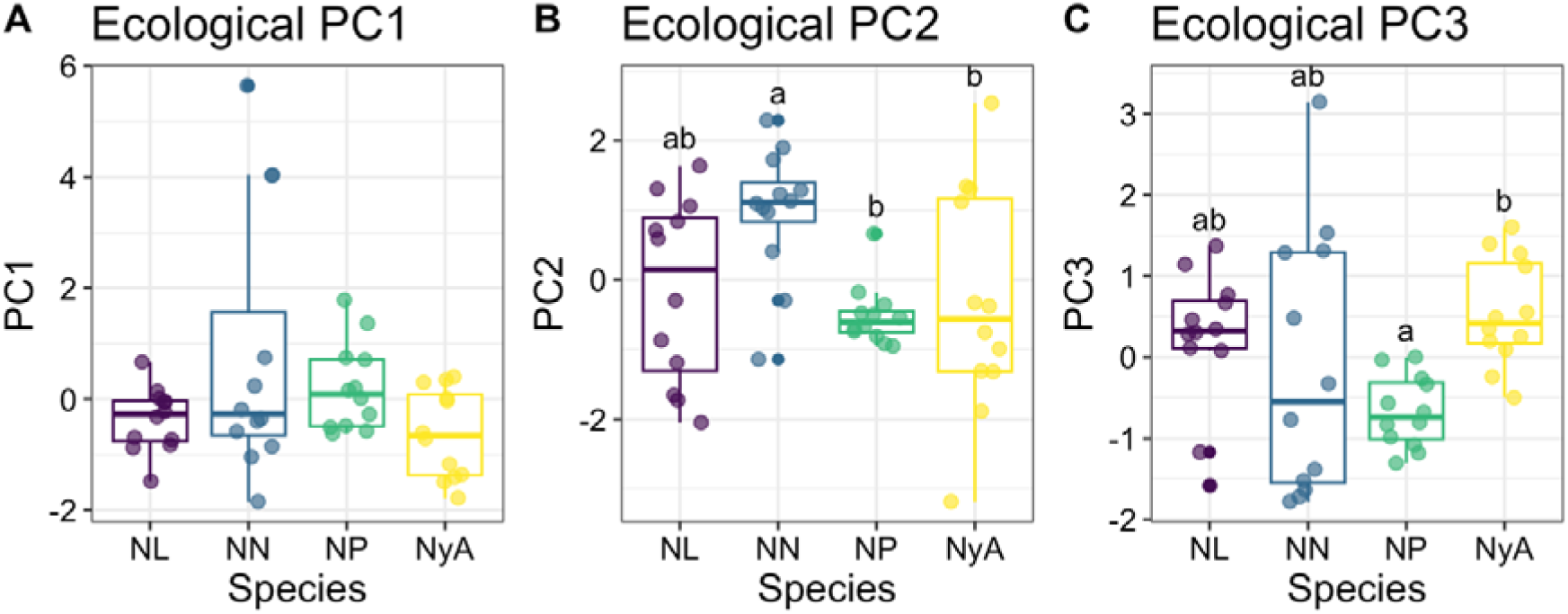
Loadings of the first three ecological PCA axes for the four investigated species, assessed with linear mixed models. Different letters indicate significant differences between species (significance level <0.05). NL=*N. lutea*, NN=*N. nucifera*, NP=*N. peltata*, NyA=*N. alba*.

### 3.2 Functional niches

For the functional niche determination, the first three PCA axes were used, which accounted for 72.1% of the trait variation (Fig. 5A-B-C). PC1 (43.2% explained variance) was mainly related to leaf size and structure. Positive values of PC1 were associated to high leaf area and carbon content, as well as proportion of petiole dry weight, leaf dry matter content and total chlorophylls, though with lower loadings (Table 1). Negative values of PC1 were instead associated to high specific leaf area. PC2 (18.9% explained variance) was related to leaf resource-use efficiency and was negatively correlated with phosphorus and specific leaf area. PC3 (9.9% explained variance), on the other hand, reflected a gradient of high to low leaf dry matter content.

**Figure 5.**
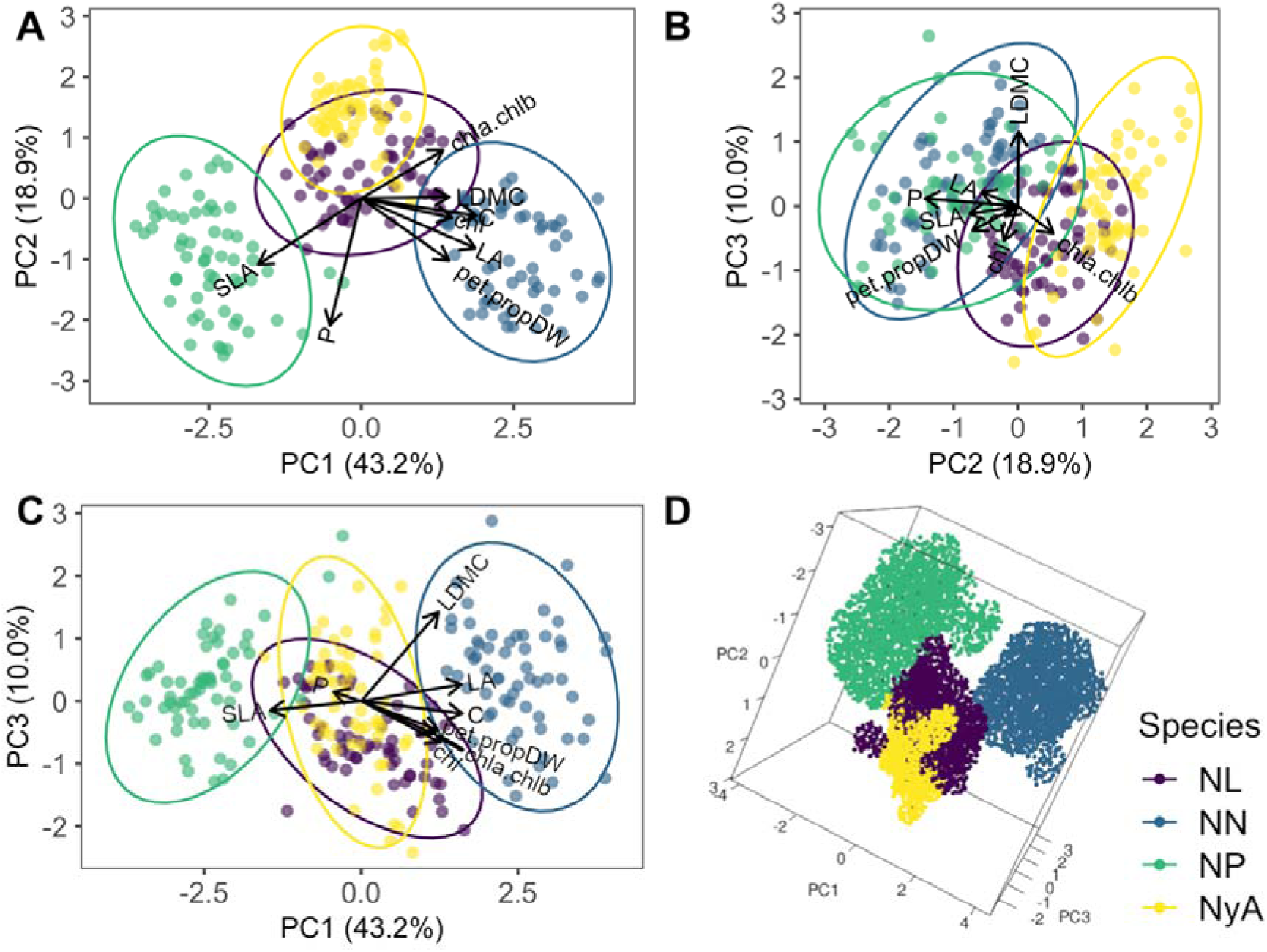
First three axes of functional PCA (A-B-C) and 3-dimensional representation of the functional niche (D). Percentage of explained variance of each axis is indicated in brackets. NL = *N. lutea*, NN = *N. nucifera*, NP = *N. peltata*, NyA = *N. alba*. For full traits names see Table 1.

The biggest mean relative niche size was found for *N. nucifera* (36.8%) followed by *N. peltata* (34.2%), *N. lutea* (18.3%) and *N. alba* (15.8%). The uncertainty estimates showed that *N. alba* has a significantly smaller relative niche size compared to *N. nucifera* and *N. peltata*, while no significant difference in niche size was observed for any other species pair (Fig. 3C, Supporting Information, Heading 4). Compared to the ecological niches, functional niches of the investigated species were more distinct: *N. nucifera* had 100.0% mean niche uniqueness, followed by *N. peltata* (99.8%), *N. lutea* (72.4%) and *N. alba* (68.1%). All estimates of niche uniqueness were significantly >0, and *N. nucifera* and *N. peltata* had significantly higher uniqueness compared to *N. alba* and *N. lutea* (Fig. 3D, Supporting Information, Heading 4). *N. nucifera* had a distinct niche that occupies the functional space described by large leaf size (indicative also of high chlorophylls content per fresh weight, as the two variables were correlated), high proportion of petiole dry weight and high phosphorus content (Fig. 5D). At the other edge of PC1, with higher specific leaf area and leaf nutrients content, and lower leaf size, laid *N. peltata*. Between these two species there were *N. lutea* and *N. alba*, with intermediate traits values. This distribution along the first PCA axis is supported by the linear mixed models (Fig. 6A, Supplementary Information, Heading 5). *N. alba* overlaps significantly with *N. lutea* functional niche (16.8% of all species volume, Supplementary Information, Heading 4), but it differentiates by a lower leaf P (and N) content (Fig. 5), although no significant difference between the two species’ PCA loadings was observed for any of the three axes considered (Fig. 6). The functional niches of *N. peltata* and *N. nucifera*, on the other hand, are not significantly overlapping with any of the investigated species. In fact, they present significantly different loadings of PC1 and PC2 axes compared to *N. lutea* and *N. alba*: in PC1, as described above, *N. peltata* and *N. nucifera* occupy opposite edges, while in PC2 they both occupy the negative edge of the axis, with high leaf phosphorus and nitrogen content (Fig. 6).

**Figure 6.**
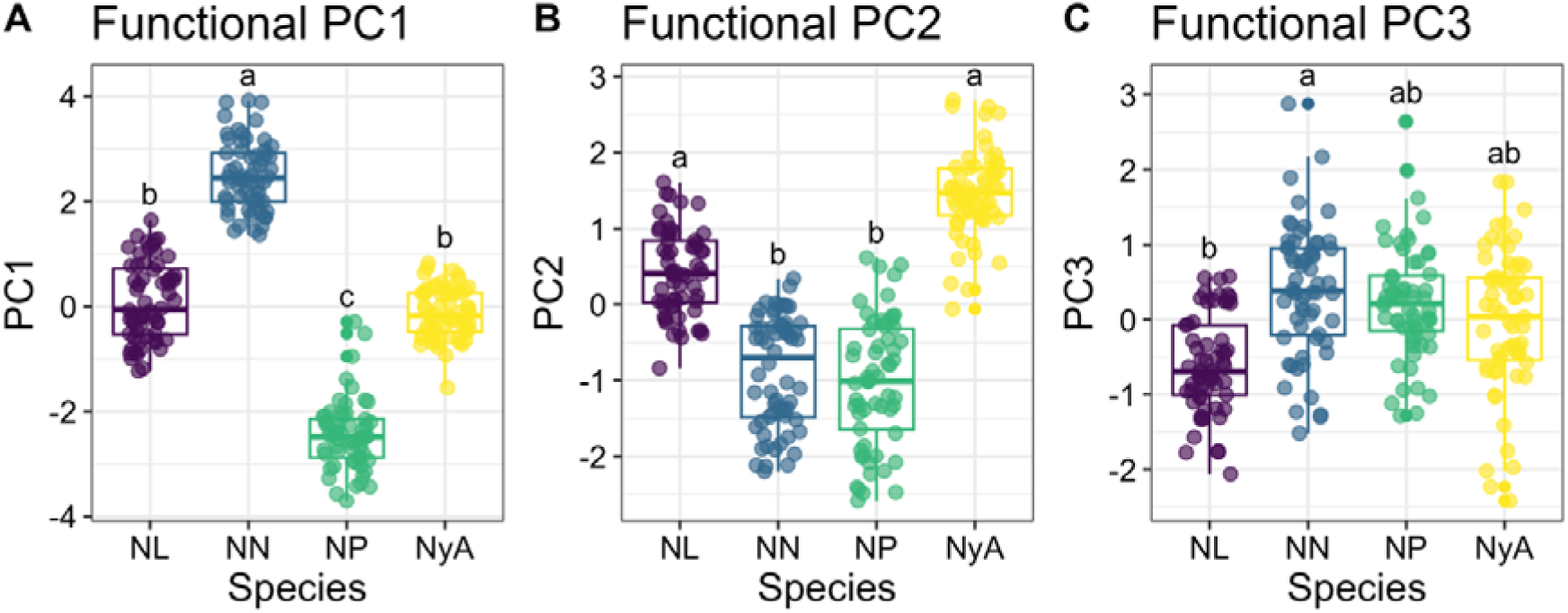
Loadings of the first three functional PCA axes for the four investigated species, assessed with linear mixed models. Different letters indicate significant differences between species (significance level <0.05). NL=*N. lutea*, NN=*N. nucifera*, NP=*N. peltata*, NyA=*N. alba*.

Based on our evidence, the functional space occupied by floating-leaved macrophytes is primarily bounded by the niches of *N. nucifera* and *N. peltata*. *N. nucifera* is the species that shows the most extreme values of leaf functional traits, and thus the greatest plasticity, presenting the highest minimum and maximum values of leaf area (196220 to 702858 mm^2^), proportion of petiole dry weight (40.22 to 68.79%), leaf dry matter content (99.3 to 242.8 mg g^−1^) and total chlorophylls (22.81 to 67.7 μg cm^−2^), and the lowest values of proportion of petiole area (3.89 to 12.33%) and specific leaf area (4.81 to 9.36 mm^2^ mg^−1^), compared to the other species. On the other side of the functional space, *N. peltata* had quite opposite behavior, presenting the lowest minimum and maximum values of leaf area (6793 to 22370 mm^2^), total chlorophylls (16.04 to 39.15 μg cm^−2^) and carbon content (40.22 to 42.73%), and the highest of specific leaf area (8.60 to 25.46 mm^2^ mg^−1^) and phosphorus content (0.25 to 0.73%).

## 4. DISCUSSION

This paper provides a robust, comprehensive perspective on the ecological and functional niches of four key floating-leaved macrophytes (*N. nucifera*, *N. alba*, *N. peltata*, and *N. lutea*). They share much of their ecological niche, suggesting occupation of fairly similar environments, but they clearly differ functionally, indicating differentiation in functional response to similar environmental conditions, and probably a differential use of resources. From an ecological point of view, this evidence is severely limited by the fact that – in the area studied, and more in general in overexploited lowlands – the habitats potentially colonized by macrophytes show a general, common status of very poor quality (Bolpagni & Piotti 2016, Bolpagni et al. 2020). Nevertheless, the four species considered here show a peculiar functional connotation that reinforces the available knowledge about their distribution patterns and adaptive abilities to perturbations, suggesting the existence of predefined functional thresholds. For instance, the functional peculiarities of each species deeply influence the temporal dynamics and spatial structure of communities and habitats they occupy. In the invaded areas of northern Italy, *N. nucifera* showed significant enhanced photosynthetic efficiency compared with native hydrophytes (Tóth et al. 2019). The species itself has mostly emergent leaves, allowing for greater water column-atmosphere interaction, including larger under-canopy light availability, than the other three nymphaeid species. All of this has the potential to trigger important cascading effects – both biogeochemical and biological – on the functioning of colonized environments (Ribaudo et al. 2012; Pannard et al. 2024), thus, there is a clear need for further investigation related to the ecosystem implications of functional traits.

### 4.1 General functional findings

Only *N. nucifera* presented a rather wide ecological niche, mainly explained by its presence in sites with extremely high conductivity values. On the other hand, *N. peltata* shows a smaller niche size, indicative of a more specialist behavior, distinguished in part from other species by its affinity for habitats with higher sediment density. This is typical of reclamation canals that are periodically dug for irrigation, where *N. peltata* retains substantial stands (Piccoli & Gerdol 1983), as it finds growing conditions similar to the natural streams no longer present in the northern Italian landscape. All species have functional niches of similar size, yet both *N. nucifera* and *N. peltata* have very distinctive functional spaces, entirely separated from the other species. This differentiation is due to the leaf size, but not only that. Indeed, both these species have a high leaf nutrients content (nitrogen and phosphorus), but also a different contribution of specific leaf area, carbon and petiole. Besides, the high chlorophyll content (per fresh weight) of *N. nucifera*, also contributes to its niche distinction, as it is collinear with leaf area. Interestingly, *N. alba* and *N. lutea* have a smaller functional niche size, occupying an intermediate portion of the “floating-leaved macrophytes” functional space, coupled with a relatively large ecological niche. This would suggest a lower ability to adapt to different environmental conditions (e.g. showing less functional plasticity). The opposite can be said for *N. peltata*, which shows a small ecological niche relatively to its functional niche size, suggesting high plasticity enclosed in a small ecological space. Our hypotheses were partially corroborated, as only *N. alba* showed smaller niche size and high overlap, while *N. peltata* presented high niche uniqueness, and the invasive *N. nucifera* had a highly unique functional niche.

### 4.2 Insights into species-specific niches

*N. lutea* shows intermediate features in terms of both ecological and functional niche. This confirms its reputation as a top-competitor species in aquatic environments (Bornette et al. 2008; Paillisson & Marion 2011). Previous studies have shown that *N. lutea* is able to modulate its response and resource-use strategies to variable conditions and can inhabit even highly eutrophic environments (Dalla Vecchia & Bolpagni, 2022; Dalla Vecchia et al. 2024). Thus, the functional plasticity of this species is triggered by changes in the environment, allowing it to increase its performance under favorable conditions (Kordyum & Klimenko 2013; Nowak et al. 2015), or to adapt to a more conservative strategy under harsh conditions (Richards et al. 2011; Klok & van der Velde 2017). However, although *N. lutea* populations can occupy a wide range of environmental conditions, with a range of electrical conductivity of nearly 400 µS cm^−1^ and nitrate concentration as high as 9.5 mg L^−1^ (Dalla Vecchia et al. 2024), based on our results, it has not yet established under extreme environmental conditions, represented by very high conductivity and nutrients content where only *N. nucifera* was found.

Similarly to *N. lutea*, *N. alba* also occupies intermediate portions of the ecological and functional “floating-leaved macrophytes” space, largely overlapping with *N. lutea*. SzaLkowski & Kłosowski (1999) and Pełechaty (2007) also observed similar and broad environmental tolerance or requirements for both species. Klok & van der Velde (2017), moreover, observed a comparable functional response of these species to variation in environmental drivers. However, adding the functional niche information, we see that the functional niche of *N. alba* is slightly shifted compared to that of *N. lutea*, towards a less acquisitive side of the functional spectrum. In fact, although *N. alba* has similar PCA loadings as *N. lutea*, it has lower leaf nutrients content (phosphorus and nitrogen), indicative of a more conservative resource-use strategy and lower competitiveness (Ding et al. 2019). Indeed, the overall decline in *N. alba* populations has been attributed to hybridization with introduced ornamental varieties (Nierbauer et al. 2014) and habitat degradation (Parveen et al. 2022). However, habitat degradation is also occurring for *N. lutea*, as they share similar environmental conditions, although the latter species does not show the same trend of decline. Therefore, we suggest that the limited plasticity given by the small functional niche coupled with the lower leaf efficiency represented by the lower nutrients content may extend the evidence based on environmental conditions, and partly explain the higher susceptibility of *N. alba* populations to degradation of aquatic ecosystems.

*N. peltata*, on the other hand, shows its specificities in both ecological and functional niches. First, this species does not exhibit the same variability in environmental tolerance, and the size of its ecological niche narrows mainly due to specific sediment quality requirements. In fact, we found *N. peltata* mainly in lowland habitats where sediments were poor in organic matter content and had a higher density due to the abundance of fine clay particles, opposite to the sediment features of sites colonized by *N. nucifera*. SzaLkowski & Kłosowski (1999) also observed a separate ecological niche of *N. peltata* compared to other nymphaeid species, distinguished by a higher mineral content in sediments. In contrast to the specificity of its ecological niche, which potentially limits the spread of this species, we observed a large and unique functional niche occupying the most acquisitive corner of the “floating-leaved macrophytes” functional space. Indeed, *N. peltata* is distinguished from other species not only by the small size and high nutrients content (also typical of *N. nucifera*) of its leaves, but also by a simple leaf structure (defined by high specific leaf area and low carbon content). This is in line with previous findings that described *N. peltata*’s high nitrogen and phosphorus content in biomass, stored in the root system and transferred to photosynthetic organs during the growing season (Brock et al. 1983b), thus confirming the active role of this species in the nutrient cycling of aquatic habitats. Markovskaya and coauthors (2019) also observed high specific leaf area in *N. peltata*, a feature that increases the photosynthetic efficiency of its leaves. The relatively low proportion of petiole dry weight observed in our study further aligns with the overall acquisitive strategy of this species, because it indicates a lower relative biomass allocation in petioles (not as photosynthetically active as leaf blades), compared to other nymphaeid species. In addition, *N. peltata* commonly reproduces *via* seeds and produces persistent seed banks (Smits et al. 1990; Wang et al. 2005), behavior that could increase the genetic diversity of *N. peltata* populations, promoting greater traits variability despite limited environmental ranges (Liao et al. 2013).

Finally, *N. nucifera* represents the only alien and invasive species in our study. Our results confirm the competitiveness and invasive potential of this species, given by a very large environmental niche combined with a unique functional niche. Its ecological niche crosses most niches of the other European nymphaeids, spreading along axes of water quality variability, extending to extreme values of conductivity (greater than 1500 µS cm^−1^), only excluding sediments with high density and low organic matter content. Overall, this indicates a marked generalist behavior. However, our hypothesis of functional niche similarity with native species was not supported as it was separated from the other species studied. This is in contrast with findings by Dalle Fratte et al. (2019), who did not observe differentiation of *N. nucifera* from other nymphaeids. This discrepancy is likely due to the fact that Dalle Fratte and coauthors (2019) represented the species groups based on the CSR strategy (*sensu* Grime & Pierce 2012), calculated using leaf area, leaf dry matter content and specific leaf area, whereas here we expanded the number of traits to represent a more comprehensive functional niche, including pigments which had a primary role in niche distinction. In this study, the uniqueness of *N. nucifera*’s niche was mainly given by leaf structure, in terms of large size and dry matter content, as well as proportion of petiole dry weight, both traits indicating a more conservative strategy, and by high nitrogen and phosphorus content in leaves. Also, the higher leaf chlorophylls concentration indicates high photosynthetic efficiency. It seems that *N. nucifera* combines acquisitive and conservative traits, and still manages to show high invasive potential. In one of our study sites, the Mantua lakes system, it was observed to rapidly colonize the available space and outcompete native nymphaeid species (*N. alba*, *N. lutea*) remarkably effectively (Pinardi et al. 2021). One reason for its ability to survive and thrive even in highly organic sediments might be its sophisticated aeration system, based on a two-way gas flow, which helps to keep its submerged organs oxygenated even under anoxic conditions (Große 1996). Besides, the extremely large leaf area of *N. nucifera* is a strong indicator of high competitiveness (Díaz et al. 2016), which allows it to effectively shade other species and rapidly gain space by forming monospecific stands. Indeed, competitive traits are typical of invasive species (Guo et al. 2022). However, large leaf area may come at a cost: large emerging leaf blades need strong support, which could explain the high proportion of petiole dry weight and leaf dry matter content observed in our study, again highlighting the importance of petioles in influencing resource-use strategies in nymphaeids (Dalla Vecchia & Bolpagni 2022).

## 5. CONCLUSIONS

This study demonstrated how the investigated nymphaeid species, while sharing more or less similar environmental conditions, are strongly differentiated with respect to their functional niche, showing adaptations not only morphological (e.g. leaf size) but also biochemical (e.g. leaf efficiency). Overall, our data strengthened the main known drivers for functional variation in floating-leaved rooted macrophytes, namely water conductivity and sediment quality, quantifying clear ecological thresholds.

The hypervolume approach made it possible to highlight aspects of species behavior, from plasticity and specialization to competitiveness. Particularly, three functional plant types have been identified: i) highly acquisitive leaves with high specific leaf area, low leaf dry matter content and high nutrients content (*N. peltata*); ii) acquisitive leaves with large leaf size and high chlorophyll and nutrients content, although with conservative traits such as high leaf dry matter content and low specific leaf area (*N. nucifera*); iii) and leaves with intermediate functional traits (*N. lutea* and *N. alba*). Indeed, the identification of different functional strategies implies a differentiation of effects and roles of species and communities in the aquatic environment. In this sense, a dynamic investigation is needed, including eco-physiological traits and seasonality of responses in future studies.

## AUTHORS CONTRIBUTION STATEMENT

ADV and RB worked together to conceive and develop the ideas. MBC and MMA helped collecting the data. ADV took the lead in analyzing the data and writing the manuscript. All authors contributed critically to the drafts and gave final approval for publication.

## DATA AVAILABILITY STATEMENT

Data will be provided on accessible repositories (e.g. Dryad or Try-database).

## ACKNOWLEDGEMENTS

We especially thank Dr. Alex Laini (University of Turin, IT) for his fundamental active contribution to the design and development of statistical analyses, and the entire macroDIVERSITY project consortium, Dr. Paolo Villa and Erika Piaser (IREA-CNR, IT), prof. Andrea Coppi and Dr. Lorenzo Lastrucci (University of Florence, IT). We also thank the students Davide Taglialatela, Mickey Cucit, Beatrice Cattabiani, Matteo Amoruso, Beatrice Fois and Elisa Marinozzi for their help in laboratory activities. The authors declare no conflicts of interest.

## FUNDING INFORMATION

This work was supported by the project “macroDIVERSITY”, funded by the Ministry of Education, University and Research, PRIN 2017 [grant number 2017CTH94H]. ADV has benefited from the equipment and framework of the COMP-HUB Initiative, funded by the ‘Departments of Excellence’ program of the Italian Ministry for Education, University and Research (MIUR, 2018-2022). This work has also benefited from the equipment and framework of the COMP-R Initiative, funded by the ‘Departments of Excellence’ program of the Italian Ministry for University and Research (MUR, 2023-2027).

## Notes

### Competing Interest Statement

The authors have declared no competing interest.

