## Supplementary Information for "Ecological and functional niches comparison reveals differentiated resource-use strategies and ecological thresholds in four key floating-leaved macrophytes"

^1^Department of Chemistry, Life Sciences and Environmental Sustainability, University of Parma, Parma (Italy), ^2^Institute for Electromagnetic Sensing of the Environment, National Research Council of Italy (CNR-IREA), Milan (Italy), ^3^Department of Biology, University of Florence, Florence (Italy), ^4^ Department of Design, Technology Architecture, Land and Environment, University of Rome La Sapienza, Rome (Italy)

**SI1:** Overview of environmental variables of sampled study sites, including type of habitat, presence of species and mean (standard deviation in brackets) of environmental variables. If no standard deviation is provided, only one population was sampled in that study site. NL=*Nuphar lutea*; NN=*Nelumbo nucifera*; NP=*Nymphoides peltata*; NyA=*Nymphaea alba*; SPC=electrical conductivity; DIC=dissolved inorganic carbon; DRSi=dissolved reactive silica; SRP=soluble reactive phosphorus; sed.OM=sediment organic matter content; sed.TP=sediment total phosphorus; sed.dens=sediment density.

| **Site** | **Habitat**  **type** | **Species** | **X** | **Y** | **Depth** m | **pH** | **SPC** μS/cm | **Nitrate** mg/L | **DRSi** μg/L | **DIC** mE/L | **SRP** μg/L | **sed.OM** % | **sed.TP** μg/g | **sed.dens** g/mL |
| --- | --- | --- | --- | --- | --- | --- | --- | --- | --- | --- | --- | --- | --- | --- |
| Annone | Lake | NL, NyA | 9.4° | 45.8° | 1.5 (0.28) | 7.73 (0.47) | 295 (2.83) | 0.08 (0.01) | 468.7 (110.34) | 2.41 (0.58) | <4 (0) | 23.05 (2.47) | 894.65 (265.09) | 1.1 (0.01) |
| Case Baldi | Channel | NP | 12.1° | 44.8° | 0.4 | 7.24 | 818.3 | 1.5 | 1201.84 | 4.38 | 18.7 | 1.18 | 452 | 1.76 |
| Cei | Lake | NyA | 11.0° | 45.9° | 1.3 | 7.8 | 271 | 0.04 | 395.42 | 2.45 | 0 | 34.31 | 2628.98 | 1.05 |
| Chiusi | Lake | NL | 12.0° | 43.1° | 1.3 (0.6) | 8.19 (0.48) | 566.33 (23.12) | 0.05 (0.09) | 831.16 (40.16) | 3.04 (0.35) | 5.86 (5.28) | 18.4 (17.07) | 561.17 (278.13) | 1.23 (0.17) |
| Codigoro | Channel | NP | 12.1° | 44.8° | 0.65 | 7.45 | 917.6 | 0.73 | 1038.1 | 3.41 | 21.48 | 4.31 | 630.02 | 1.36 |
| Comabbio | Lake | NN, NyA | 8.7° | 45.8° | 0.8 (0.57) | 8.52 (0.17) | 260.5 (6.36) | 0.06 (0.01) | 2259.03 (56.53) | 1.81 (0.28) | <4 (0) | 33.15 (42.56) | 623.12 (508.05) | 1.33 (0.41) |
| Ferrara | Channels | NP | 11.9° | 45.0° | 1.02 (0.29) | 7.87 (0.36) | 538.17 (172.42) | 1.35 (0.95) | 1225.6 (233.28) | 2.36 (0.87) | 10.39 (7.67) | 3.11 (1.22) | 520.88 (169.05) | 1.64 (0.12) |
| Fimon | Lake | NyA | 11.5° | 45.5° | 0.73 (0.25) | 7.5 (0.91) | 259.33 (3.51) | 0.03 (0.04) | 621.96 (50.61) | 2.12 (0.71) | <4 (0.41) | 29.67 (8.2) | 887.15 (313.41) | 1.06 (0.06) |
| Gorro | Lake | NyA | 9.9° | 44.5° | 1.5 | 8.05 | 517.9 | 0.04 | 1189.03 | 4.59 | 9.78 | 3.93 | 150.33 | 1.63 |
| Mantova | Lakes | NL, NN | 10.7° | 45.2° | 1.16 (0.71) | 7.72 (0.3) | 402.88 (17.44) | 7 (2.45) | 1032.82 (118.65) | 3.35 (0.59) | 46.39 (26.24) | 16.62 (3.01) | 1493.36 (491.31) | 1.21 (0.29) |
| Massaciuccoli | Channel | NN | 10.4° | 43.8° | 0.7 | 8 | 2214 | 0.15 | 1908.46 | 4.41 | 251.7 | 44.1 | 1117.3 | 0.92 |
| Massarosa | Pond | NN | 10.3° | 43.9° | 0.15 (0.07) | 7.44 (0.06) | 1327 (26.87) | 0.12 (0.02) | 3421.19 (1587) | 6.64 (0.01) | 340.54 (204.71) | 18.35 (19.71) | 634.74 (399.56) | 1.29 (0.29) |
| Monticchio | Lake | NyA | 15.6° | 40.9° | 2 (0.28) | 8.91 (0.02) | 458.5 (0.71) | 0.05 (0.04) | 210.13 (24.98) | 3.34 (0.1) | <4 (0.99) | 28.06 (6.78) | 753.25 (138.19) | 1.04 (0.02) |
| Pusiano | Lake | NL, NyA | 9.3° | 45.8° | 1.43 (0.06) | 8.41 (0.11) | 217.33 (2.52) | 0.84 (0.45) | 70.94 (13.07) | 6.34 (3.83) | 9.43 (5.91) | 14.07 (2.84) | 498.97 (310.24) | 1.16 (0.02) |
| Torbiere del  Sebino | Swamp | NL | 10.0° | 45.6° | 0.77 (0.58) | 9.57 (0.13) | 290.67 (5.13) | 2.05 (3.11) | 1124.04 (245.13) | 4.24 (1.17) | 42.4 (7.04) | 15.37 (14.51) | 747.13 (156.87) | 1.42 (0.51) |
| Vallazza | Swamp | NL, NP,  NyA | 10.8° | 45.1° | 1.2 (0.08) | 7.97 (0.16) | 389.64 (44.5) | 1.55 (0.7) | 951.35 (436.33) | 3.4 (0.28) | 25.62 (22.22) | 16.89 (7.84) | 1352.76 (427) | 1.24 (0.15) |
| Varese | Lake | NN | 8.7° | 45.8° | 0.32 (0.04) | 8.46 (0.43) | 267 (9.9) | 0.03 (0.03) | 304.95 (22.35) | 3.42 (0.9) | 3.81 (5.39) | 25.46 (5.57) | 1177.97 (116.15) | 1.14 (0.06) |

**SI2:** Pearson’s correlation tables of environmental variables and functional traits. For labels meaning refer to SI2; sed.dens=sediment density; chl.a.g, chl.b.g, car.g=chlorophyll-a, chlorophyll-b, carotenoids per fresh weight; chl.a.cm, chl.b.cm, car.cm=chlorophyll-a, chlorophyll-b, carotenoids per cm^2^; chla.chlb=chlorophyll-a/chlorophyll-b ratio; chlab.car=chlorophylls/carotenoids ratio.

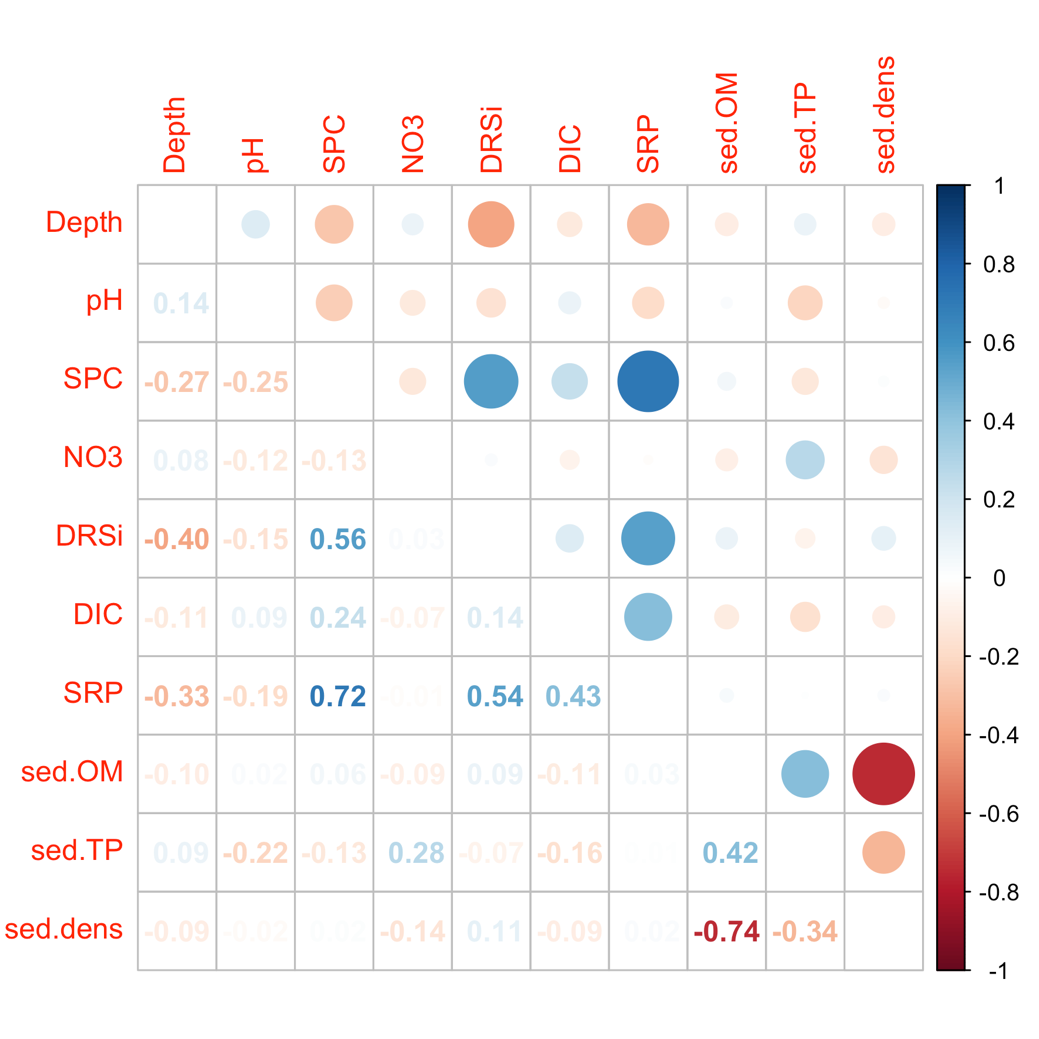

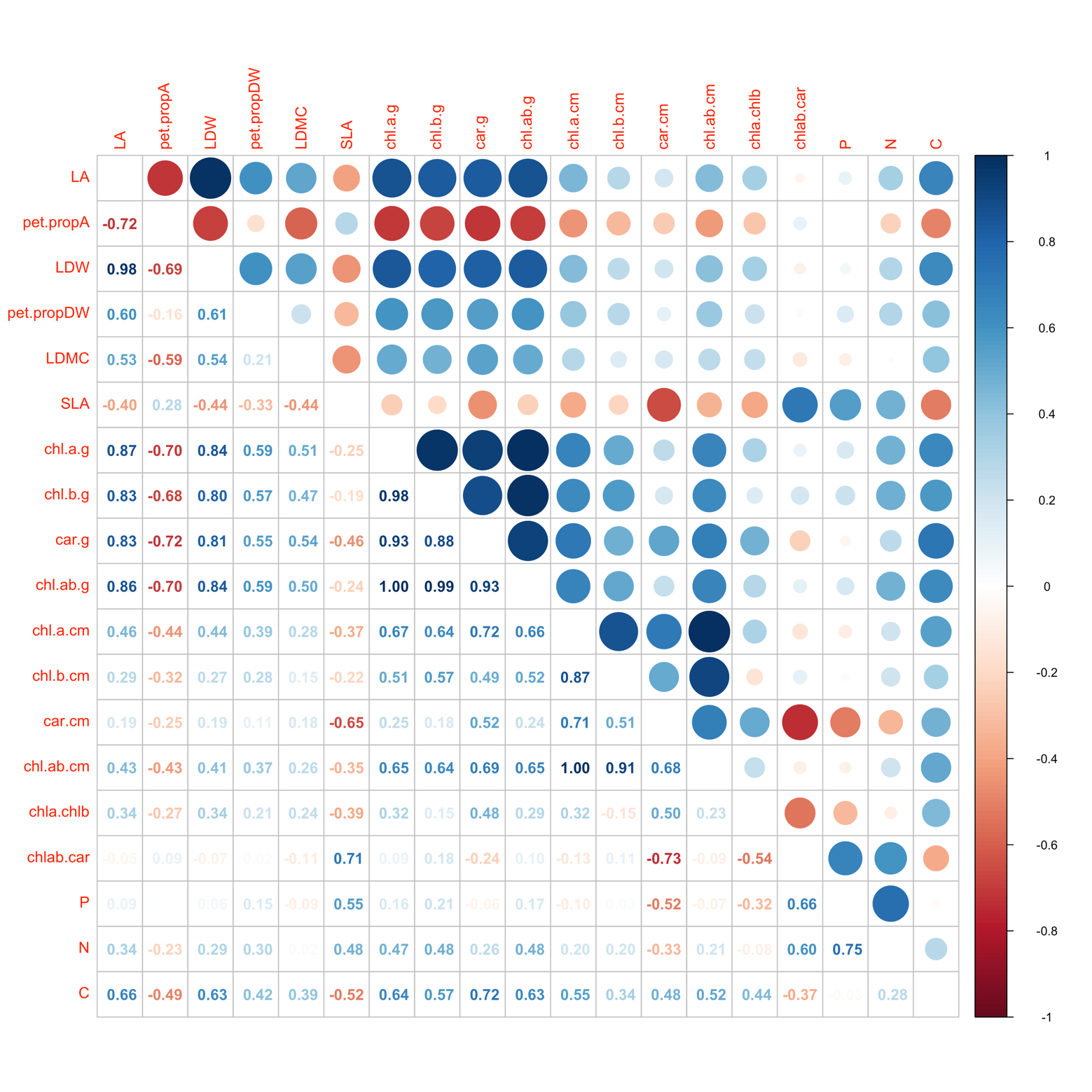

**SI3**: post-hoc pairwise comparisons between species in environmental variables and functional traits. A summary table is provided indicating mean (and standard deviations in brackets) of traits and environmental variables for each species, where significant differences are indicated by different letters and highlighted in bold. Parametric or non-parametric test and post-hoc test choice are indicated. Significance level set at 0.05. “diff”=mean difference between species; “lwr”=lowest difference; “upr”=highest difference; “p adj”=adjusted p-value; LMM=linear mixed models; NL=*N.r lutea*; NN=*N. nucifera*; NP=*N. peltata*; NyA=*N. alba*; SPC=electrical conductivity; DIC=dissolved inorganic carbon; DRSi=dissolved reactive silica; SRP=soluble reactive phosphorus; sed.OM=sediment organic matter content; sed.TP=sediment total phosphorus; sed.density=sediment density; LA=leaf area; pet.propA=proportion of petiole area; LDW=leaf dry weight; pet.propDW=proportion of petiole dry weight; LDMC=leaf dry matter content; SLA=specific leaf area; chlab=total chlorophylls; P, N and C=leaf phosphorus, nitrogen and carbon content.

| **Variable** |  | ***Nuphar***  ***lutea*** | ***Nelumbo***  ***nucifera*** | ***Nymphoides***  ***peltata*** | ***Nymphaea***  ***alba*** |
| --- | --- | --- | --- | --- | --- |
| Depth | m | 1.03 (0.52) | 0.87 (0.72) | 0.98 (0.3) | 1.33 (0.46) |
| pH |  | 8.41 (0.78) | 7.86 (0.48) | 7.82 (0.35) | 8.11 (0.64) |
| SPC | µS cm^-2^ | 390.8 (123.5) | 674.9 (614.9) | 535.4 (211.4) | 326.8 (110.5) |
| DIC | mE L^-1^ | 3.49 (0.76) | 3.87 (1.53) | 2.93 (0.89) | 3.58 (2.47) |
| Nitrate | mg L^-1^ | **2.26 (3.16) ab** | **3.37(3.89) ab** | **1.37(0.85) a** | **0.26 (0.45) b** |
| DRSi | µg L^-1^ | **900.0 (376.7) ab** | **1482.6 (1161.2) a** | **1038.1 (313.5) ab** | **686.0 (674.9) b** |
| SRP | µg L^-1^ | **24.0 (20.8) ab** | **104.1 (143.6) a** | **12.4 (7.2) ab** | **8.0 (16.3) b** |
| sed.OM | % | **18.7 (10.5) a** | **24.6 (15.8) a** | **5.7 (4.4) b** | **21.5 (11.5) a** |
| sed.TP | µg g^-1^ | 984.7 (459.4) | 1260.4 (534.6) | 700.6 (319.8) | 868.5 (719.3) |
| sed.density | g mL^-1^ | **1.23 (0.26) b** | **1.18 (0.27) b** | **1.53 (0.20) a** | **1.18 (0.22) b** |
| LA | mm^2^ | **75610 (19339) b** | **417330 (115826) a** | **14934 (3722) c** | **74119 (24912) b** |
| pet.propA | % | **0.20 (0.05) a** | **0.08 (0.02) b** | **0.21 (0.06) a** | **0.18 (0.03) a** |
| LDW | g | **10.6 (3.8) b** | **58.4 (18.4) a** | **0.9 (0.3) c** | **10.9 (4.4) b** |
| pet.propDW | % | **0.44 (0.07) b** | **0.53 (0.07) a** | **0.38 (0.08) bc** | **0.38 (0.06) c** |
| LDMC | mg g^-1^ | **110.7 (15.2) b** | **162.3 (29.3) a** | **109.3 (24.6) b** | **121.9 (24.7) b** |
| SLA | mm^2^ mg^-1^ | **7.58 (1.59) b** | **7.3 (1.1) b** | **18.0 (4.0) a** | **7.0 (1.3) b** |
| chlab | µg g^-1^ | **905.5 (372.7) b** | **2640.5 (634.2) a** | **806.4 (308.3) b** | **798.2 (368.5) b** |
| chlab | µg cm^-2^ | **37.4 (10.6) ab** | **41.3 (9.8) a** | **26.0 (5.4) c** | **30.4 (10.3) bc** |
| P | % | **0.28 (0.09) b** | **0.37 (0.11) ab** | **0.48 (0.13) a** | **0.17 (0.04) c** |
| N | % | **2.97 (0.51) b** | **3.64 (0.56) a** | **3.61 (0.52) a** | **2.20 (0.33) c** |
| C | % | **42.85 (0.50) b** | **43.84 (1.13) a** | **41.31 (0.92) c** | **42.48 (0.43) b** |

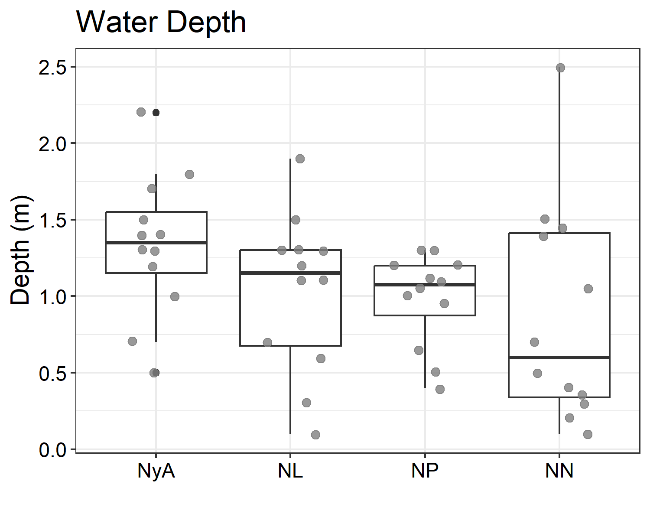

**Depth** (ANOVA + Tukey)

|  | diff | lwr | upr | p adj |
| --- | --- | --- | --- | --- |
| NN-NL | -0.163 | -0.733 | 0.408 | 0.8715 |
| NP-NL | -0.053 | -0.623 | 0.518 | 0.9947 |
| NyA-NL | 0.300 | -0.270 | 0.870 | 0.5031 |
| NP-NN | 0.110 | -0.460 | 0.680 | 0.9550 |
| NyA-NN | 0.463 | -0.108 | 1.033 | 0.1488 |
| NyA-NP | 0.353 | -0.218 | 0.923 | 0.3616 |

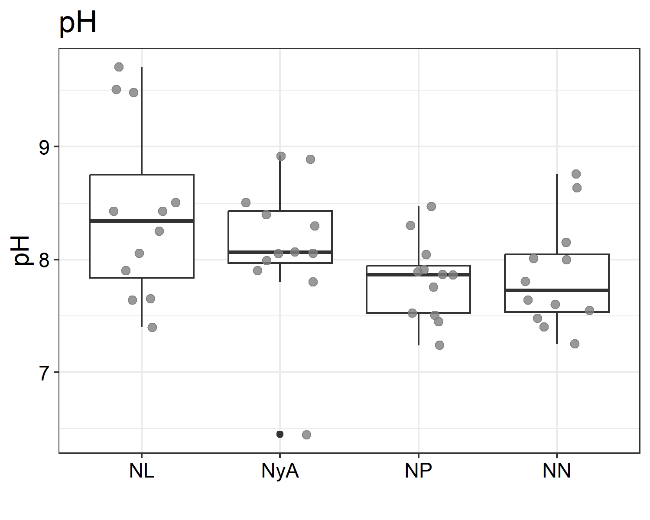

**pH** (ANOVA + Tukey)

|  | diff | lwr | upr | p adj |
| --- | --- | --- | --- | --- |
| NN-NL | -0.557 | -1.193 | 0.079 | 0.1052 |
| NP-NL | -0.594 | -1.230 | 0.042 | 0.0748 |
| NyA-NL | -0.303 | -0.939 | 0.334 | 0.5867 |
| NP-NN | -0.038 | -0.674 | 0.599 | 0.9986 |
| NyA-NN | 0.254 | -0.382 | 0.890 | 0.7112 |
| NyA-NP | 0.292 | -0.344 | 0.928 | 0.6150 |

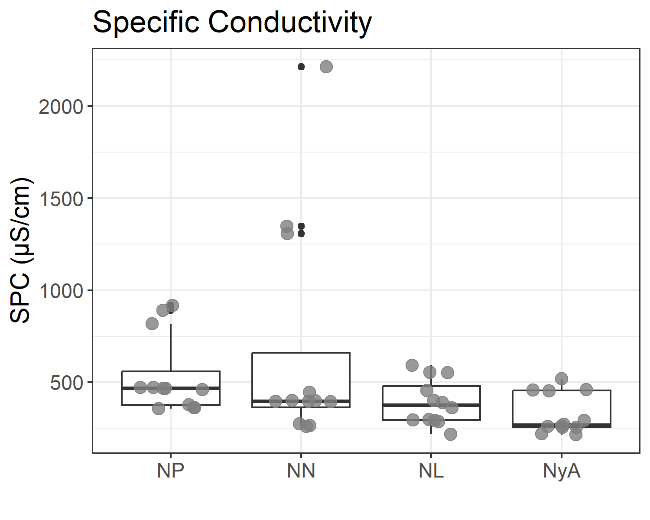

**Electrical conductivity** (ANOVA + Tukey)

|  | diff | lwr | upr | p adj |
| --- | --- | --- | --- | --- |
| NN-NL | 284.1 | -81.6 | 649.8 | 0.1775 |
| NP-NL | 144.5 | -221.2 | 510.2 | 0.7181 |
| NyA-NL | -64.0 | -429.7 | 301.7 | 0.9658 |
| NP-NN | -139.6 | -505.2 | 226.1 | 0.7393 |
| NyA-NN | -348.1 | -713.8 | 17.6 | 0.0673 |
| NyA-NP | -208.5 | -574.2 | 157.1 | 0.4330 |

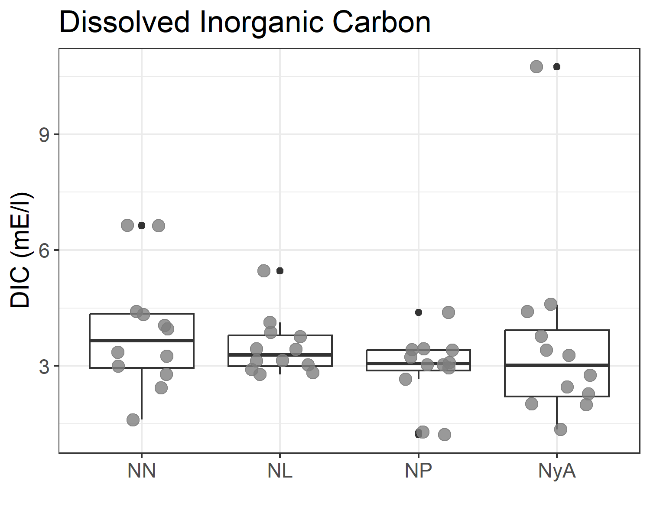
**Dissolved inorganic carbon** (ANOVA +

Tukey)

|  | diff | lwr | upr | p adj |
| --- | --- | --- | --- | --- |
| NN-NL | 0.382 | -1.322 | 2.085 | 0.9320 |
| NP-NL | -0.563 | -2.266 | 1.141 | 0.8143 |
| NyA-NL | 0.097 | -1.607 | 1.800 | 0.9987 |
| NP-NN | -0.944 | -2.647 | 0.759 | 0.4579 |
| NyA-NN | -0.285 | -1.988 | 1.418 | 0.9699 |
| NyA-NP | 0.659 | -1.044 | 2.362 | 0.7310 |

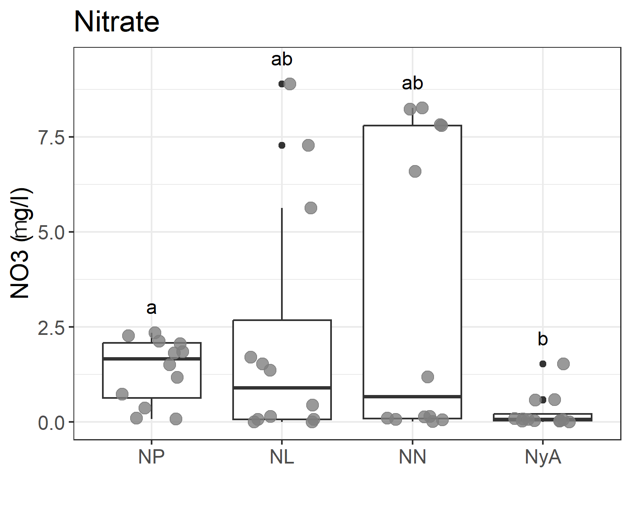
**Nitrate** (Kruskal-Wallis + Dunn)

|  | diff | p adj |
| --- | --- | --- |
| NL-NN | -0.510 | 1.0000 |
| NL-NP | -0.941 | 1.0000 |
| NN-NP | -0.430 | 0.6670 |
| NL-NyA | 1.976 | 0.1925 |
| NN-NyA | 2.487 | 0.0645 |
| NP-NyA | 2.917 | 0.0212 |

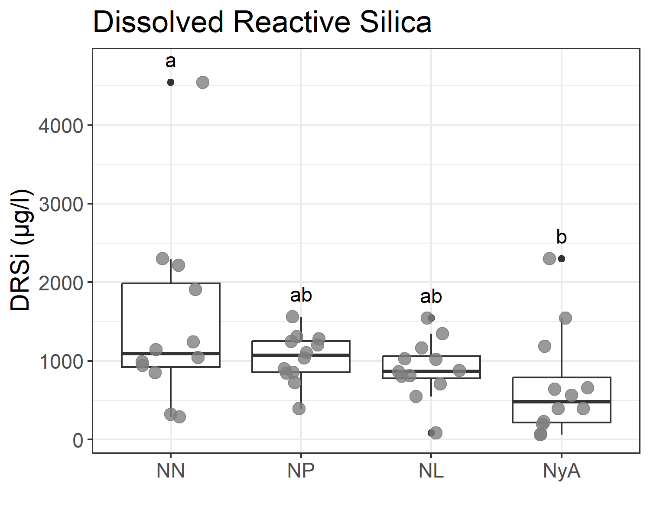

**Dissolved reactive silica** (ANOVA + Tukey)

|  | diff | lwr | upr | p adj |
| --- | --- | --- | --- | --- |
| NN-NL | 582.6 | -196.6 | 1361.8 | 0.2049 |
| NP-NL | 138.0 | -641.1 | 917.2 | 0.9646 |
| NyA-NL | -214.0 | -993.2 | 565.2 | 0.8832 |
| NP-NN | -444.6 | -1223.8 | 334.6 | 0.4326 |
| NyA-NN | -796.6 | -1575.8 | -17.4 | 0.0434 |
| NyA-NP | -352.0 | -1131.2 | 427.2 | 0.6262 |

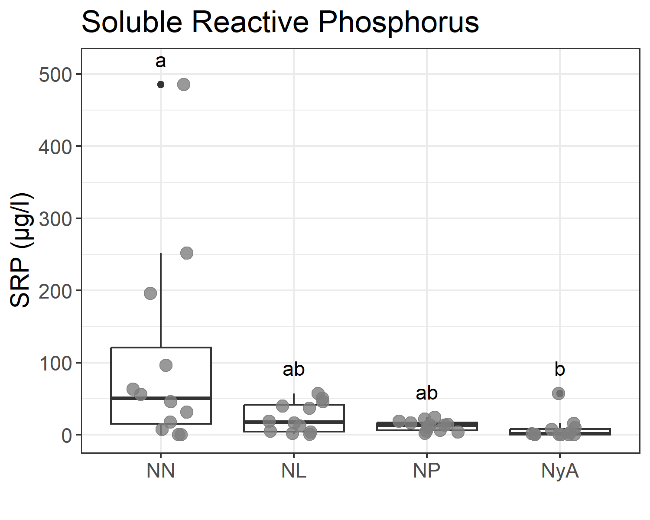

**Soluble reactive phosphorus** (Kruskal-Wallis +

Dunn)

|  | diff | p adj |
| --- | --- | --- |
| NL-NN | -1.080 | 0.5601 |
| NL-NP | 0.650 | 0.5160 |
| NN-NP | 1.730 | 0.2510 |
| NL-NyA | 2.387 | 0.0850 |
| NN-NyA | 3.467 | 0.0032 |
| NP-NyA | 1.737 | 0.3295 |

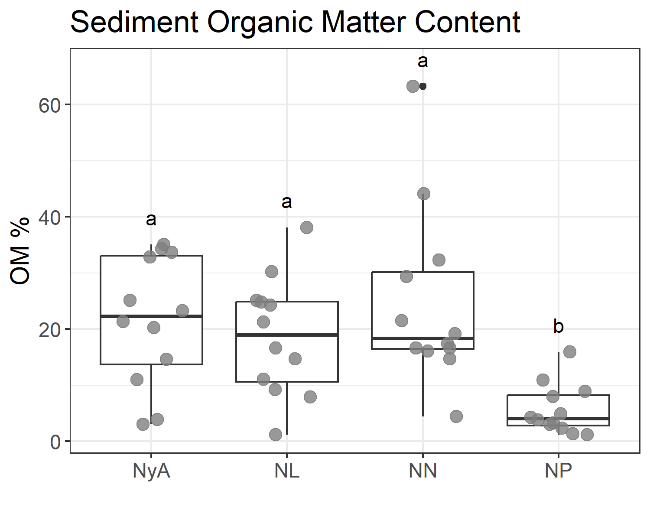
**Sediment organic matter content** (ANOVA +

Tukey)

|  | diff | lwr | upr | p adj |
| --- | --- | --- | --- | --- |
| NN-NL | 5.92 | -6.42 | 18.26 | 0.5796 |
| NP-NL | -13.05 | -25.39 | -0.72 | 0.0345 |
| NyA-NL | 2.83 | -9.51 | 15.17 | 0.9277 |
| NP-NN | -18.97 | -31.31 | -6.63 | 0.0010 |
| NyA-NN | -3.09 | -15.43 | 9.25 | 0.9082 |
| NyA-NP | 15.88 | 3.54 | 28.22 | 0.0069 |

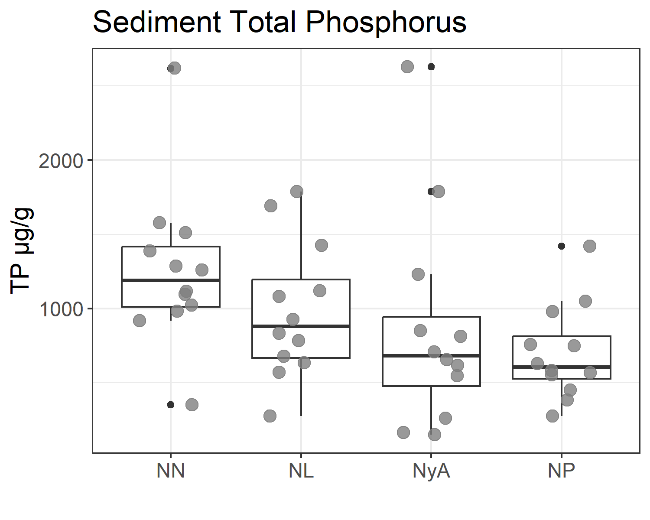

**Sediment total phosphorus** (ANOVA + Tukey)

|  | diff | lwr | upr | p adj |
| --- | --- | --- | --- | --- |
| NN-NL | 275.7 | -300.1 | 851.6 | 0.5813 |
| NP-NL | -284.1 | -859.9 | 291.8 | 0.5573 |
| NyA-NL | -116.2 | -692.1 | 459.7 | 0.9490 |
| NP-NN | -559.8 | -1135.7 | 16.1 | 0.0595 |
| NyA-NN | -392.0 | -967.9 | 183.9 | 0.2790 |
| NyA-NP | 167.8 | -408.1 | 743.7 | 0.8640 |

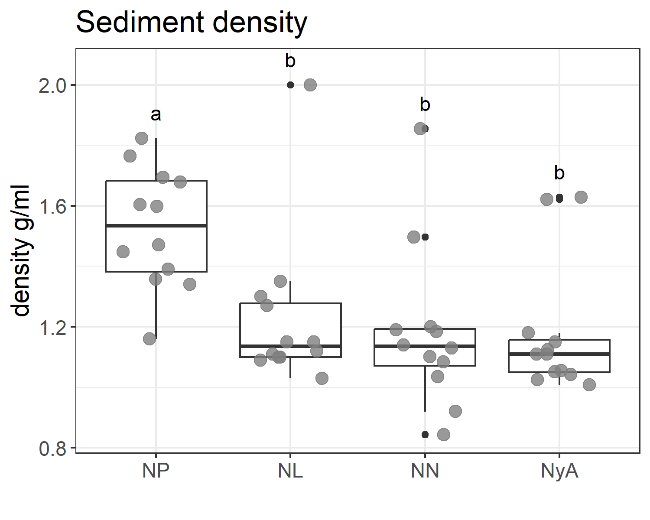

**Sediment density** (ANOVA + Tukey)

|  | diff | lwr | upr | p adj |
| --- | --- | --- | --- | --- |
| NN-NL | -0.0494 | -0.3076 | 0.2089 | 0.9562 |
| NP-NL | 0.2965 | 0.0382 | 0.5547 | 0.0188 |
| NyA-NL | -0.0554 | -0.3137 | 0.2029 | 0.9397 |
| NP-NN | 0.3458 | 0.0876 | 0.6041 | 0.0046 |
| NyA-NN | -0.0060 | -0.2643 | 0.2522 | 0.9999 |
| NyA-NP | -0.3519 | -0.6101 | -0.0936 | 0.0039 |

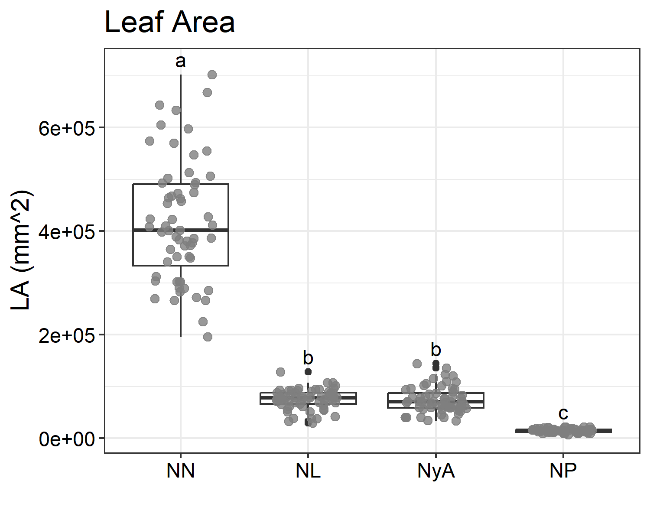
**Leaf area** (LMM + Tukey)

|  | diff | p.value |
| --- | --- | --- |
| NL-NN | -341720 | 0.0000 |
| NL-NP | 60676 | 0.0245 |
| NL-NyA | 1632 | 0.9998 |
| NN-NP | 402396 | 0.0000 |
| NN-NyA | 343351 | 0.0000 |
| NP-NyA | -59044 | 0.0298 |

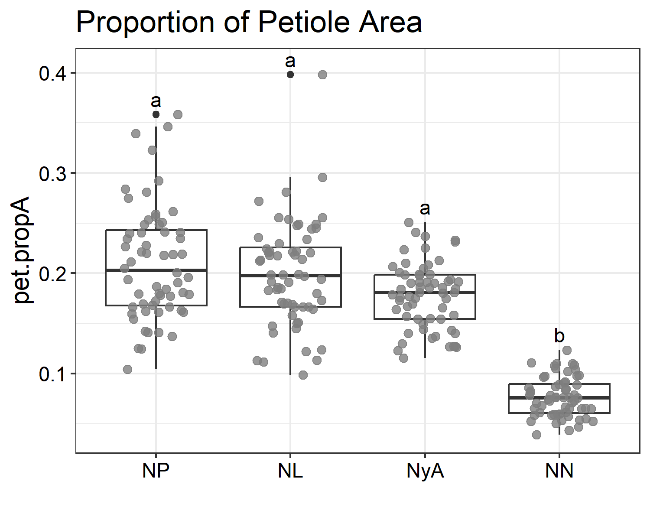

**Proportion of petiole area** (LMM + Tukey)

|  | diff | p.value |
| --- | --- | --- |
| NL-NN | 0.955 | 0.0000 |
| NL-NP | -0.054 | 0.9070 |
| NL-NyA | 0.097 | 0.6212 |
| NN-NP | -1.009 | 0.0000 |
| NN-NyA | -0.857 | 0.0000 |
| NP-NyA | 0.151 | 0.2484 |

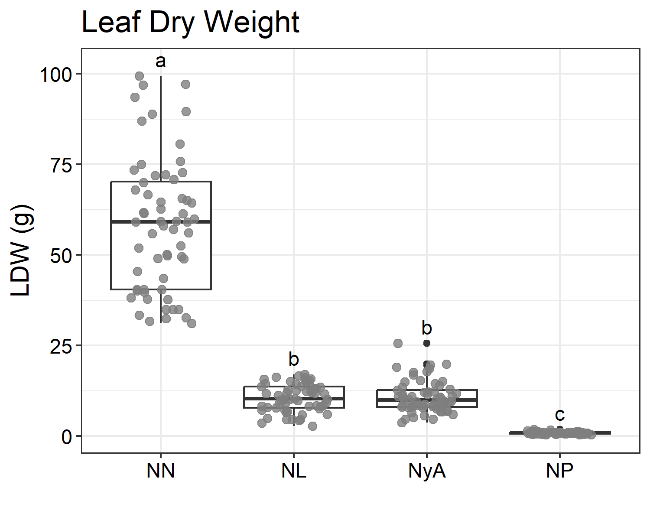

**Leaf dry weight** (LMM + Tukey)

|  | diff | p.value |
| --- | --- | --- |
| NL-NN | -1.735 | 0.0000 |
| NL-NP | 2.472 | 0.0000 |
| NL-NyA | -0.025 | 0.9978 |
| NN-NP | 4.207 | 0.0000 |
| NN-NyA | 1.710 | 0.0000 |
| NP-NyA | -2.497 | 0.0000 |

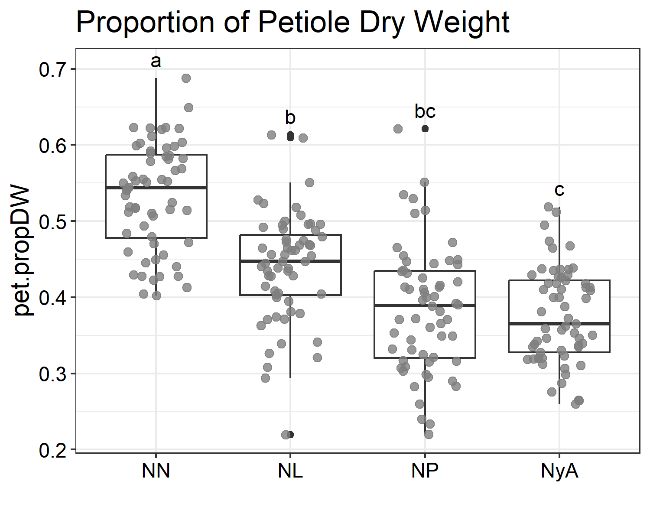

**Proportion of petiole dry weight** (LMM +

Tukey)

|  | diff | p.value |
| --- | --- | --- |
| NL-NN | -0.0934 | 0.0004 |
| NL-NP | 0.0549 | 0.0588 |
| NL-NyA | 0.0636 | 0.0214 |
| NN-NP | 0.1483 | 0.0000 |
| NN-NyA | 0.1570 | 0.0000 |
| NP-NyA | 0.0087 | 0.9763 |

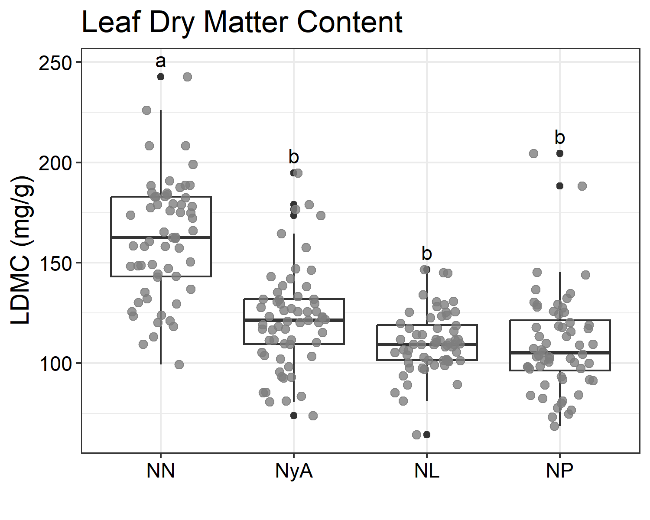
**Leaf dry matter content** (LMM + Tukey)

|  | diff | p.value |
| --- | --- | --- |
| NL-NN | -0.3760 | 0.0000 |
| NL-NP | 0.0248 | 0.9762 |
| NL-NyA | -0.0864 | 0.4834 |
| NN-NP | 0.4007 | 0.0000 |
| NN-NyA | 0.2896 | 0.0001 |
| NP-NyA | -0.1112 | 0.2646 |

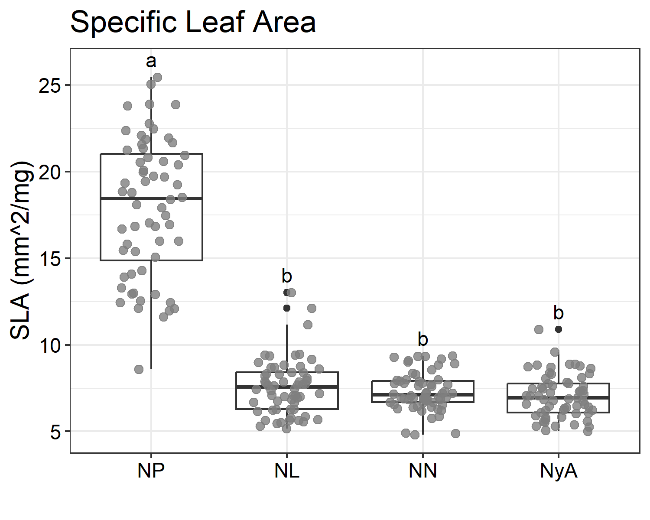

**Specific leaf area** (LMM + Tukey)

|  | diff | p.value |
| --- | --- | --- |
| NL-NN | 0.0280 | 0.9718 |
| NL-NP | -0.8562 | 0.0000 |
| NL-NyA | 0.0692 | 0.7041 |
| NN-NP | -0.8842 | 0.0000 |
| NN-NyA | 0.0412 | 0.9178 |
| NP-NyA | 0.9253 | 0.0000 |

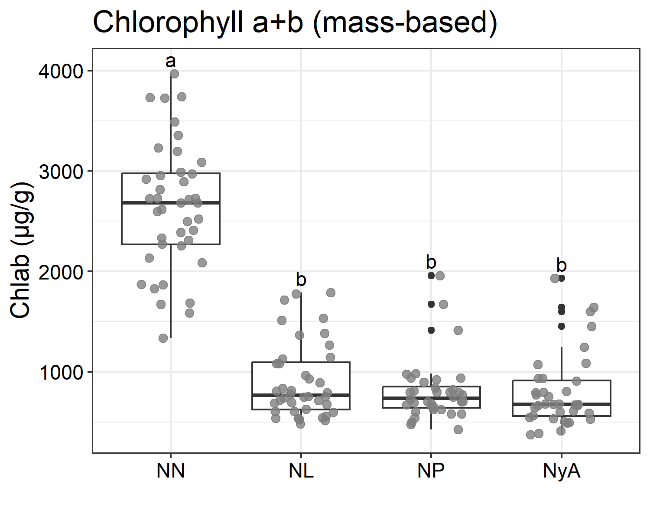

**Chlorophylls** *a*+*b*, fresh weight based (LMM +

Tukey)

|  | diff | p.value |
| --- | --- | --- |
| NL-NN | -1.115 | 0.0000 |
| NL-NP | 0.092 | 0.8515 |
| NL-NyA | 0.133 | 0.6472 |
| NN-NP | 1.207 | 0.0000 |
| NN-NyA | 1.248 | 0.0000 |
| NP-NyA | 0.042 | 0.9831 |

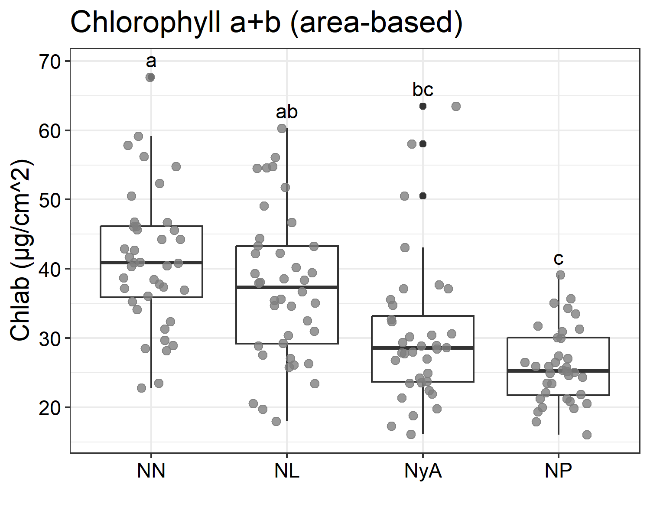

**Chlorophylls** *a*+*b*, area based (LMM + Tukey)

|  | diff | p.value |
| --- | --- | --- |
| NL-NN | -0.125 | 0.4729 |
| NL-NP | 0.341 | 0.0015 |
| NL-NyA | 0.211 | 0.0818 |
| NN-NP | 0.466 | 0.0000 |
| NN-NyA | 0.336 | 0.0018 |
| NP-NyA | -0.130 | 0.4462 |

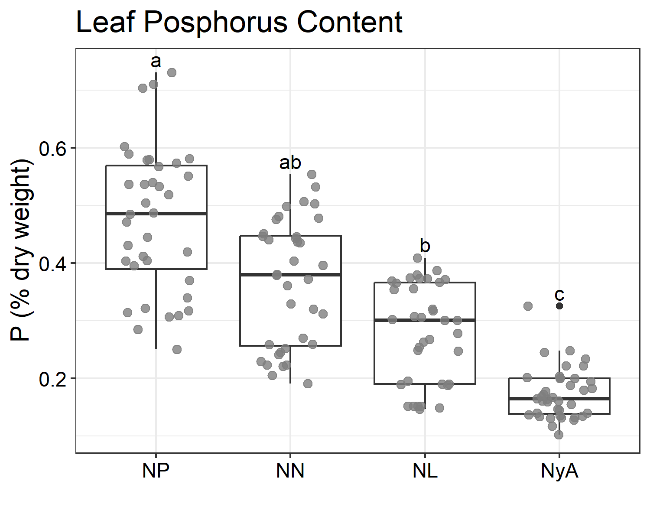
**Leaf phosphorus content** (LMM + Tukey)

|  | diff | p.value |
| --- | --- | --- |
| NL-NN | -0.277 | 0.0980 |
| NL-NP | -0.558 | 0.0001 |
| NL-NyA | 0.444 | 0.0025 |
| NN-NP | -0.281 | 0.0931 |
| NN-NyA | 0.721 | 0.0000 |
| NP-NyA | 1.002 | 0.0000 |

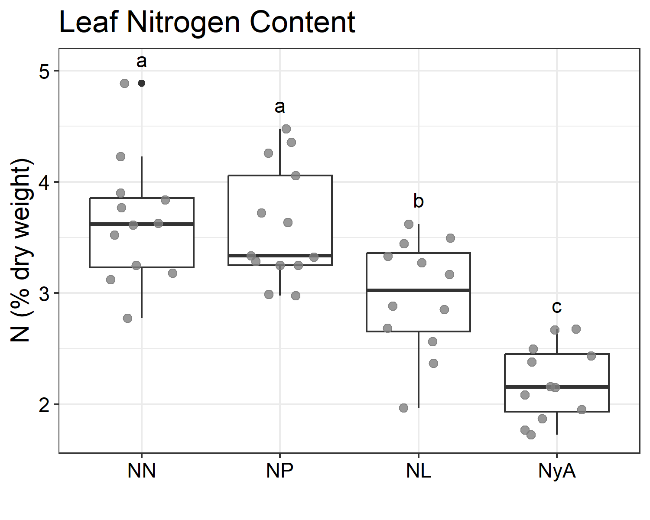

**Leaf nitrogen content** (ANOVA + Tukey)

|  | diff | lwr | upr | p adj |
| --- | --- | --- | --- | --- |
| NN-NL | 0.672 | 0.141 | 1.203 | 0.0081 |
| NP-NL | 0.638 | 0.117 | 1.159 | 0.0108 |
| NyA-NL | -0.774 | -1.305 | -0.242 | 0.0018 |
| NP-NN | -0.034 | -0.555 | 0.487 | 0.9981 |
| NyA-NN | -1.445 | -1.977 | -0.914 | 0.0000 |
| NyA-NP | -1.412 | -1.932 | -0.891 | 0.0000 |

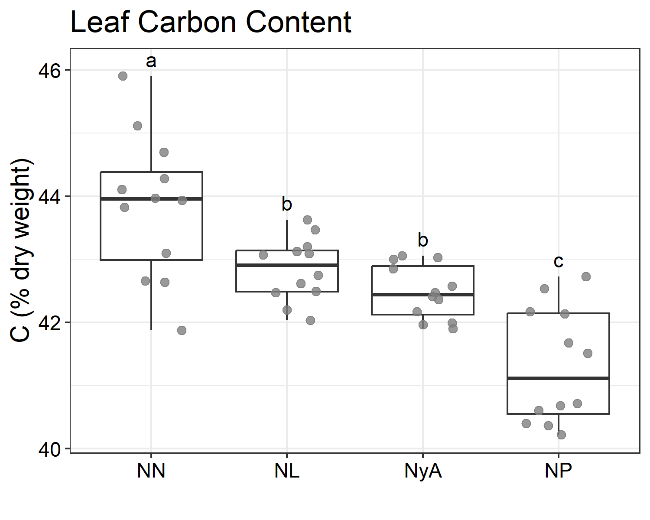

**Leaf carbon content** (ANOVA + Tukey)

|  | diff | lwr | upr | p adj |
| --- | --- | --- | --- | --- |
| NN-NL | 0.998 | 0.127 | 1.869 | 0.0191 |
| NP-NL | -1.534 | -2.405 | -0.662 | 0.0001 |
| NyA-NL | -0.363 | -1.234 | 0.509 | 0.6843 |
| NP-NN | -2.532 | -3.403 | -1.660 | 0.0000 |
| NyA-NN | -1.361 | -2.232 | -0.489 | 0.0008 |
| NyA-NP | 1.171 | 0.299 | 2.042 | 0.0045 |

**SI4**: Uncertainty estimates of hypervolumes size, uniqueness, and intersection obtained with bootstrapping on 199 permutated hypervolumes on each species. Relative niche size is the proportion of niche occupied by each species relative to the total niche occupied by all species. Relative niche uniqueness is the proportion of unique niche of each species relative to its total niche volume. Relative niche intersection (overlap) is the proportion of niche shared by two species compared to the union of all volumes. Mean values, as well as2.5 and 97.5 quantiles (Q 2.5 and Q 97.5 respectively), are provided. NL=*N. lutea*, NN=*N. nucifera*, NP=*N. peltata*, NyA=*N. alba*.

|  |  | ECOLOGICAL NICHE | | | FUNCTIONAL NICHE | | |
| --- | --- | --- | --- | --- | --- | --- | --- |
|  | Species | Mean | Q 2.5 | Q 97.5 | Mean | Q 2.5 | Q 97.5 |
| Relative niche size | NL | 0.2439 | 0.1455 | 0.0257 | 0.1826 | 0.1134 | 0.2793 |
|  | NN | 0.7589 | 0.2027 | 0.0097 | 0.3675 | 0.2675 | 0.4574 |
|  | NP | 0.0558 | 0.0388 | 0.0064 | 0.3416 | 0.2526 | 0.4423 |
|  | NyA | 0.1437 | 0.1105 | 0.0035 | 0.1583 | 0.1064 | 0.2225 |
| Relative niche uniqueness | NL | 0.3471 | 0.1213 | 0.7097 | 0.7244 | 0.6229 | 0.8209 |
|  | NN | 0.8291 | 0.5915 | 0.9336 | 0.9999 | 0.9990 | 1.0000 |
|  | NP | 0.4826 | 0.1651 | 0.7758 | 0.9978 | 0.9905 | 1.0000 |
|  | NyA | 0.3791 | 0.1392 | 0.7506 | 0.6807 | 0.5162 | 0.8163 |
| Relative niche intersection | NL - NN | 0.1086 | 0.0322 | 0.2317 | 0.0000 | 0.0000 | 0.0006 |
|  | NL - NP | 0.0640 | 0.0062 | 0.1247 | 0.0005 | 0.0000 | 0.0030 |
|  | NL - NyA | 0.1777 | 0.0464 | 0.3027 | 0.1682 | 0.1036 | 0.2299 |
|  | NN - NP | 0.0131 | 0.0000 | 0.0371 | 0.0000 | 0.0000 | 0.0000 |
|  | NN - NyA | 0.0444 | 0.0000 | 0.1034 | 0.0000 | 0.0000 | 0.0000 |
|  | NP - NyA | 0.0427 | 0.0023 | 0.1030 | 0.0010 | 0.0000 | 0.0050 |

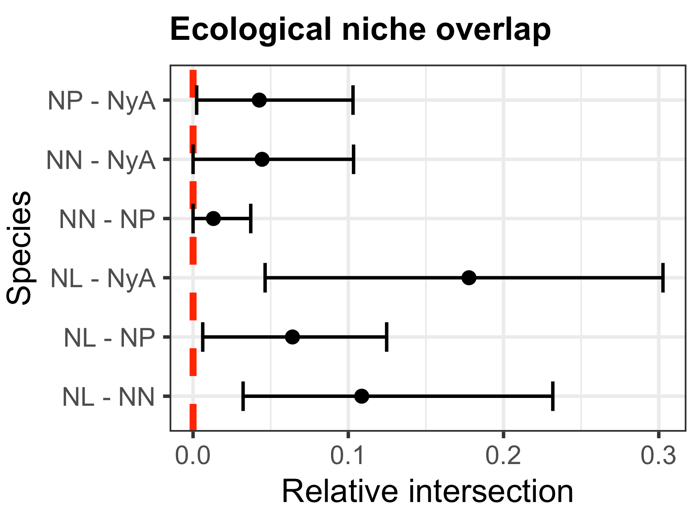

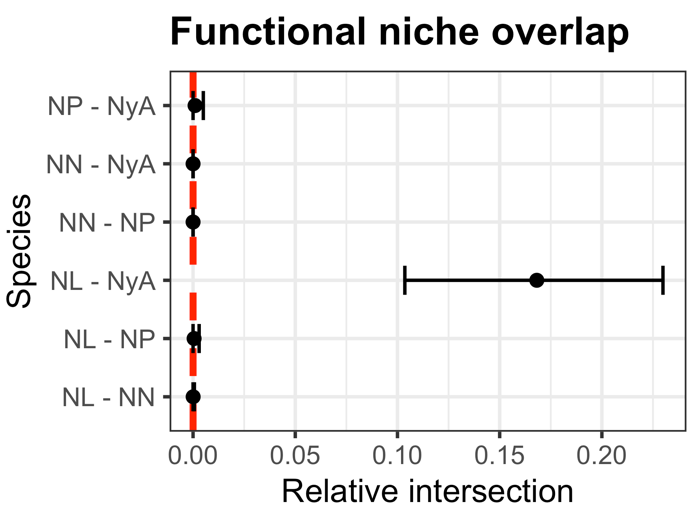

**SI5:** post-hoc pairwise comparisons between species in PCA-based linear mixed models. “diff”=mean difference between species; NL=*N. lutea*; NN=*N. nucifera*; NP=*N.s peltata*; NyA=*N. alba*.

Ecological PC1: PC1 ~ Species + (1|Year)

Ecological PC2: PC2 ~ Species + (1|Year)

Ecological PC3: PC3 ~ Species + (1|Year)

|  | **PC1** | | **PC2** | | **PC3** | |
| --- | --- | --- | --- | --- | --- | --- |
|  | diff | p-value | diff | p-value | diff | p-value |
| NL - NN | -1.031 | 0.4159 | -1.107 | 0.2689 | 0.496 | 0.7345 |
| NL - NP | -0.408 | 0.9436 | 0.378 | 0.9417 | 1.108 | 0.1805 |
| NL - NyA | 0.404 | 0.9365 | 0.184 | 0.9910 | -0.14 | 0.9923 |
| NN - NP | 0.623 | 0.6875 | **1.485** | **0.0218** | 0.613 | 0.4512 |
| NN - NyA | 1.435 | 0.0597 | **1.291** | **0.0483** | -0.636 | 0.4048 |
| NP - NyA | 0.812 | 0.4645 | -0.194 | 0.9773 | **-1.249** | **0.0179** |

Functional PC1: PC1 ~ Species + (1|Year) + (1|Population)

Functional PC2: PC2 ~ Species + (1|Year) + (1|Population)

Functional PC3: PC3 ~ Species + (1|Year) + (1|Population)

|  | **PC1** | | **PC2** | | **PC3** | |
| --- | --- | --- | --- | --- | --- | --- |
|  | diff | p-value | diff | p-value | diff | p-value |
| NL - NN | **-2.466** | **<0.0001** | **1.558** | **<0.0001** | **-0.909** | **0.0244** |
| NL - NP | **2.427** | **<0.0001** | **1.777** | **<0.0001** | -0.719 | 0.1623 |
| NL - NyA | 0.137 | 0.9586 | -0.745 | 0.0969 | -0.496 | 0.4092 |
| NN - NP | **4.893** | **<0.0001** | 0.218 | 0.8231 | 0.19 | 0.8808 |
| NN - NyA | **2.602** | **<0.0001** | **-2.303** | **<0.0001** | 0.413 | 0.3694 |
| NP - NyA | **-2.291** | **<0.0001** | **-2.521** | **<0.0001** | 0.223 | 0.8131 |
